# Extinctions in the marine plankton preceded by stabilizing selection

**DOI:** 10.1101/531947

**Authors:** Manuel F. G. Weinkauf, Fabian G. W. Bonitz, Rossana Martini, Michal Kučera

## Abstract

Unless they adapt, populations facing persistent stress are threatened by extinction. Theoretically, populations facing stress can react by either disruption (increasing trait variation and potentially generating new traits) or stabilization (decreasing trait variation). In the short term, stabilization is more economical, because it quickly transfers a large part of the population closer to a new ecological optimum. However, canalization is deleterious in the face of persistently increasing stress, because it reduces variability and thus decreases the ability to react to further changes. Understanding how natural populations react to intensifying stress reaching terminal levels is key to assessing their resilience to environmental change such as that caused by global warming. Because extinctions are hard to predict, observational data on the adaptation of populations facing extinction are rare. Here, we make use of the glacial salinity rise in the Red Sea as a natural experiment allowing us to analyse the reaction of planktonic Foraminifera to stress escalation in the geological past. We analyse morphological trait state and variation in two species across a salinity rise leading to their local extinction. One species reacted by stabilization in shape and size, detectable several thousand years prior to extinction. The second species reacted by trait divergence, but each of the two divergent populations remained stable or reacted by further stabilization. These observations indicate that the default reaction of the studied Foraminifera is canalization, and that stress escalation did not lead to the emergence of adapted forms. An inherent inability to breach the global adaptive threshold would explain why communities of Foraminifera and other marine protists reacted to Quaternary climate change by tracking their zonally shifting environments. It also means that populations of marine plankton species adapted to response by migration will be at risk of extinction when exposed to stress outside of the adaptive range.

## 1 Introduction

Extinctions are difficult to study due to their unpredictability in recent environments (Moritz and Agudo 2013). This is because it is difficult to quantify stress reactions over ecologically relevant time-scales, which cannot be simulated in the laboratory, or because the severity and ultimate outcome of the stress reaction is hard to predict.

Mechanistically, environmental stress influences population morphology by decreasing the fitness of specimens which show a high degree of developmental instability (Lens et al. 2002, Hendrickx et al. 2003). On the population level, this can lead to either stabilizing selection when a certain phenotype is the preferred survivor toward the stress (e.g. Van Valen 1965) or disruptive selection when higher variation better guarantees the survival of the population in a rapidly changing environment (e.g. Bull 1987). Both stabilizing and disruptive selection are detectable by assessing the phenotypic variation (i.e. the range of realized phenotypes, Schmalhauzen 1949) of the population, which can therefore be used as a measure for developmental stability. Shape and size of organisms have long been hypothesized to reflect the influence of environmental stress on the physiology of an organism during its lifetime (Furlow et al. 1997, Lens et al. 2002, Klingenberg 2003, Osterauer et al. 2010, Marschner et al. 2013). Therefore, a characterization of shape and size and their variation should in principle allow an assessment of the severity of stress exposure. Under this assumption, stress exposure that leads to extinction, can be expected to leave a discernible imprint on morphology in pre-extinction populations (West-Eberhard 2003).

In this regard, the sedimentary record offers a unique opportunity to study the effects of terminal stress levels (i.e. stress leading to local extinction), because here the outcome can be directly observed. This benefit comes at the cost that the sedimentary record always comprises a temporally integrated sequence, and that the environmental change that caused the local extinction can be difficult to reconstruct in some settings (compare Weinkauf et al. 2014). Shell bearing protists, such as planktonic Fo-raminifera, are perfect model systems for such studies, because they are preserved in great numbers in the fossil record (Schiebel 2002, Kučera 2007) and thus allow a robust analysis of their variation on the population level.

Here, we use two species of planktonic Foraminifera from the Red Sea sediment core KL 09 to study their morphological reaction toward terminal stress levels. Understanding the morphological reaction of Foraminifera toward environmental stress could serve as a proxy for evolvability of the present assemblages in this organismal group and possibly of the plankton at large. Past studies have shown that morphological deviations in benthic and planktonic Foraminifera can be caused by environmental forcing (Malmgren and Kennett 1976, Malmgren 1984, Baumfalk et al. 1987, Mary and Knappertsbusch 2013, Moller et al. 2013, Weinkauf et al. 2014, Knappertsbusch 2016, Brombacher et al. 2017). But since these studies quantified morphology in very different ways, the results obtained are scarce and controversial. In light of earlier results, it is reasonable to assume to see a morphological trend in the assemblage of planktonic Foraminifera associated with terminal stress levels. We hypothesize that a correlation between morphology and either environmental proxies or species abundance (as biotic indicator for the stress level the community is exposed to) exists. To test this hypothesis, we use Pleistocene sediments from the Red Sea, where several species of planktonic Foraminifera regionally disappeared from the fossil record (hereafter called local extinction) within aplanktonic zones resulting from environmental change (Fenton et al. 2000). Shortly before the onset of each aplanktonic zone, a distinct sequential local extinction pattern of different planktonic Foraminifera species can be observed, until virtually all planktonic Foraminifera are absent from the fossil record. In contrast to comparable cases where fossil material is used (e.g. Weinkauf et al. 2014, Brombacher et al. 2017), we are here in the unique situation that the extinction events can nearly exclusively be linked to salinity increase (Thunell et al. 1988, Rohling et al. 2009), allowing a qualification of the stressor which acted on the assemblage.

The study will evaluate the value of morphology as proxy for stress and the adaptive patterns acting in planktonic Foraminifera communities. By qualifying evolutionary patterns in regard to environmental stress reactions, we will be able to gain insights into the evolutionary mode acting upon planktonic Foraminifera, and to evaluate their adaptive potential to environmental change.

## 2 Material and methods

### 2.1 Sample material

For the present study we used material from piston core Geo-Tü KL 09 (19.804 ° N, 38.103 ° E, Fig. 1) taken in the Red Sea during the RV Meteor cruise M 5-2 (Nellen et al. 1996). We chose material from marine isotope stage 12 (MIS 12), specifically the range from 461.1 to 437.5 kyrs bp (age model after Grant et al. 2014), to investigate planktonic Foraminifera morphology under terminal environmental stress. The interval covers an aplanktonic zone, which occurred during the Pleistocene in the Red Sea as a result of extremely high salinities (*>*49 on the psu scale) induced by a changed circulation pattern in the Red Sea basin (Fenton et al. 2000). The MIS 12 aplanktonic zone has been chosen, because it is the most prominent and longest one preserved in KL 09. The spatial resolution for our sampling varies between 0.5 and 2 cm, for a temporal resolution of 120–500 years (higher resolution for the last 2000 years before the aplanktonic zone). Since salinity values in the Red Sea are tightly coupled with the relative sea level and *δ*^18^O values of the sea water (Thunell et al. 1988), and high resolution sea level and stable isotope reconstructions from the Red Sea exist (Rohling et al. 2009, Fig. 1), we can approximate past sea water salinity to test for its influence on foraminiferal shell morphology. While MIS 12 represents a glacial period, it was shown that the sea surface temperature in the Red Sea area rarely dropped below 24 °C (Rostek et al. 1997), which is well within the temperature tolerance levels of all species common in the Red Sea (Prell et al. 1999a, Prell et al. 1999b), so that sea surface temperature played no considerable role in environmental stress levels.

**Figure 1.**
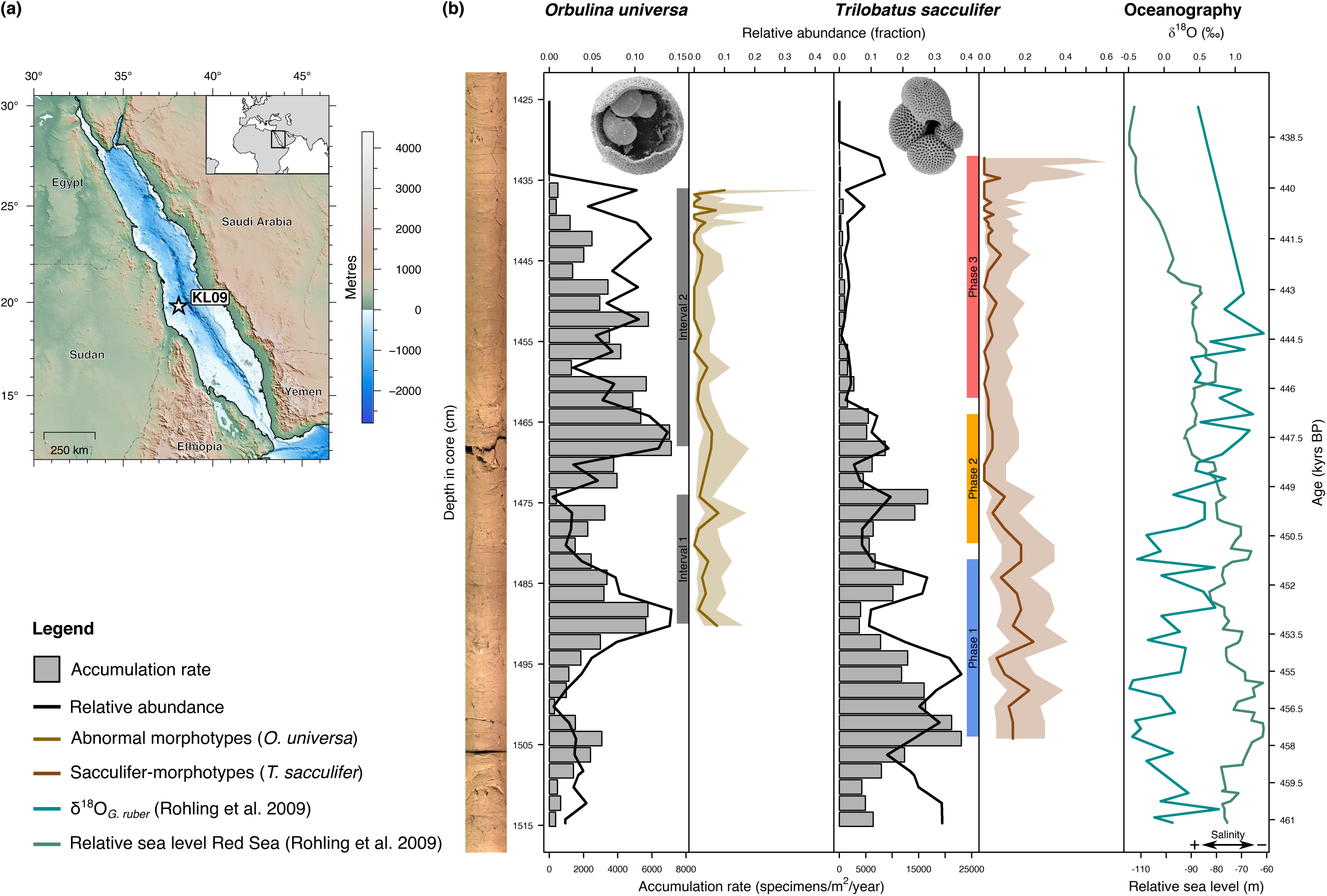
Summary of the sampling material from piston core Geo-Tü KL 09. (a) Map of the sampling area with indication of the core position. (b) Stratigraphic plot with core image. Accumulation rates, relative abundances in relation to other species of planktonic Foraminifera, and the incidence of abnormal morphotypes in *Orbulina universa* and sacculifer-morphotypes in *Trilobatus sacculifer* is indicated (shaded area depicts 95 % confidence intervals). The aplanktonic zone begins at approximately 438 kyrs BP. The two intervals of dropping abundance for *O. universa* (Intervals 1 and 2) and the three phases defined for *T. sacculifer* (Phases 1–3) based on relative abundances are indicated. The *δ*^18^O_*G ruber*_ measurements and the three-point moving average relative sea level in the Red Sea (Rohling et al. 2009) is also shown. Example scanning electron microscope images of the two investigated species are shown in the respective panels. The *O. universa* specimen was cracked open to reveal the juvenile shell inside. Image of *T. sacculifer* from Hesemann (2009).

We investigated two abundant, symbiont-bearing species of planktonic Foraminifera (Fig. 1, Suppl. S1). Both species react sensitively to salinity changes, as shown by their consistently early position within the extinction sequence of the aplanktonic zones (Fenton et al. 2000). *Orbulina universa* d’Orbigny, 1839 is characterized by a trochospiral juvenile shell that is overgrown by a spherical terminal chamber. *Trilobatus sacculifer* (Brady, 1877) Spezzaferri et al., 2015 shows a trochospiral shell, in which the terminal chamber sometimes develops a sac-like shape. Both species are surface dwellers, partly due to their symbionts’ need for light, and thus occur in comparable environments (compare Rebotim et al. 2017, Schiebel and Hemleben 2017, Meilland et al. 2019). Differences in their reactions can therefore not be the result of the exposure to considerably different environmental forcing.

### 2.2 Sample preparation and data acquisition

Sediment samples of 0.5 cm thickness were taken with a U-channel from piston core Geo-Tü KL 09, dried in an oven at 50 °C, soaked in tap-water, and washed over a 63 µm screen under flowing tap-water. The residual *>*63 µm was dried and dry-sieved over a 150 µm screen. Only the fraction *>*150 µm was used for this study to ensure that the analysed individuals would have reached their adult stage so that an ontogenetic effect on the shape analyses could be widely eliminated (Peeters et al. 1999). For census counts, only a small fraction of each sample (split with a microsplitter) has been used. Aliquots containing at least 300 specimens were investigated for their species composition and counted samples were stored without modification.

Planktonic foraminiferal specimens from representative aliquots (split with a microsplitter) were picked with a needle and transferred onto glass slides, were they were fixed in position using permanent glue. We were striving for a sample size *n* of at least 5 specimens per sample. For *O. universa* we always used the entire sample aliquot, while for *T. sacculifer* a randomly selected subsample (*n* = 50) was analysed due to the more time-consuming analysis in this species. Images for morphological analyses were taken with a Canon EOS 500D digital mirror reflex camera attached to a Zeiss Stereo.V8 binocular microscope under constant magnification.

Morphological data for *O. universa* (2774 specimens) were semi-automatically extracted from high-contrast transmitted light images using the software FIJI (ImageJ v. 1.48s, Schindelin et al. 2012). We used the shell size (Feret diameter) and shell roundness (ratio between longest and shortest axis of a fitted ellipse, 1.0 equals a sphere) (Suppl. S1), as well as the incidence of the abnormal ecophenotypes ‘*Orbulina suturalis*’ and ‘*Biorbulina bilobata*’ (Caron et al. 1987b). For *T. sacculifer* (2228 specimens), images were taken under reflected light with specimens oriented such that the apertural plane was lying horizontally, perpendicular to the direction of view (apertural standard view). In these images, 12 landmarks (Suppl. S1) were manually digitized in R v. 3.5.1 (R Core Team 2018). In 68 specimens, parts of the structures were not visible clearly enough to extract all landmarks and they were excluded from all analyses using the landmark data, leaving a dataset of 2160 specimens. Furthermore, the attribution to one of the three morphotypes of the species (trilobus, quadrilobatus, sacculifer, compare André et al. 2013) was recorded. Attribution to a morphotype was done by the same author (MFGW) for all specimens to avoid errors due to varying species concepts.

### 2.3 Morphometric data analysis

All statistical analyses have been performed in R v. 3.5.1 (R Core Team 2018). The normality of data distribution was tested with a Shapiro–Wilk test (Shapiro and Wilk 1965) and the homoscedasticity by a Fligner–Killeen test (Fligner and Killeen 1976) wherever necessary. Confidence intervals of morphological parameters were calculated via bootstrapping with the R-package ‘boot’ v. 1.3-20 (Davison and Hinkley 1997). Confidence intervals for morphotype occurrences were calculated using multinomial equations (Heslop et al. 2011). In all cases of multiple testing (e.g. pairwise tests between more than two groups), *p*-values were corrected for the false discovery rate after Benjamini and Yekutieli (2001).

To investigate trends in the incidence of abnormal morphotypes of *O. universa* and sacculifer-morphotypes of *T. sacculifer* over time, we used generalized linear models (GLM, Nelder and Wedderburn 1972) on the binomial distribution with logit as link-function. The coefficient of determination was determined according to equations by Nagelkerke (1991) (Nagelkerke-*R*^2^). For *T. sacculifer* we additionally used pairwise two-proportions *z*-tests to investigate the influence of stress levels on the incidence of sacculifer-morphotypes.

For *O. universa*, the extracted morphological parameters were subjected to traditional morphometric analyses. For *T. sacculifer* we used traditional morphometrics as well as geometric morphometric analytical methods as described in Claude (2008) and Zelditch et al. (2012). The landmark coordinates were fully Procrustes fitted using the R-package ‘shapes’ v. 1.2.4. The centroid sizes were used as size parameters for *T. sacculifer* shells. All size data were log_*e*_-transformed prior to analyses.

Traditional morphometric group-differences were investigated by a Kruskal–Wallis test (Kruskal and Wallis 1952), under circumstances followed by pairwise Mann–Whitney *U* tests (Mann and Whitney 1947). Bimodality was tested using Hartigan’s dip test (Hartigan and Hartigan 1985) as implemented in the Rpackage ‘diptest’ v. 0.75-7, and the bimodality coefficient after Ellison (1987) was calculated with the R-package ‘modes’ v. 0.7.0. Trends in variation were analysed by calculating the coefficient of variation with the 95 % confidence interval after Vangel (1996). Kendall–Theil robust line fitting model III linear regression (Kendall 1938, Theil 1950, Sen 1968) was used for all linear regression analyses. To test for correlations between size and roundness in *O. universa* we applied a Kendall rank-order correlation (Kendall 1938).

The shape of *T. sacculifer* specimens, defined as the Riemannian shape distance (Kendall 1984) of an individual landmark configuration from the grand mean shape, was investigated using geometric morphometrics. The variation of the *T. sacculifer* populations was approximated as variance of individual Riemannian shape distances within the population. Its confidence intervals and standard errors were derived by bootstrapping, and pairwise comparisons were performed by Student’s *t*-tests with adjusted degrees of freedom (number of specimens in both groups minus two, Zelditch et al. 2012). Superimposed landmark data were analysed for differences between groups using non-parametric multivariate analysis of variance (NPMANOVA, Anderson 2001) on the Euclidean distances with 999 permutation as implemented in the R-package ‘vegan’ v. 2.5-2. Ensuing pairwise comparisons used the ‘testmeanshapes’ function of the R-package ‘shapes’ v. 1.2.4 with 999 permutations. Shape changes were visualized using thin-plate splines (Bookstein 1989); and canonical variates analysis (CVA, Fisher 1936) from the R-package ‘MASS’ v. 7.3-50 (Venables and Ripley 2002) was used to analyse the shape changes between predefined groups.

We further tested all observed morphological trends against three potential models of phyletic evolution using the R-package ‘paleoTS’ v. 0.5-1 (Hunt 2006). This allows to distinguish between directional selection (general random walk), a directional pattern due to the accumulation of random change (unbiased random walk), and a system that does not change over time (stasis). The corrected Akaike information criterion (AIC_*c*_, Akaike 1974) in combination with Akaike weights (Wagenmakers and Farrell 2004) was used to decide, which model best describes the data.

## 3 Results

### 3.1 Abundance patterns

#### 3.1.1 Species abundances

*Orbulina universa* occurred at generally low abundances (5.6 % on average) that never exceeded 14.5 % of the total assemblage (Fig. 1). The species shows two abundance peaks in the studied interval, in case of the second event leading to local extinction. The first abundance peak occurred around 453 kyrs bp, and was followed by a rapid decline in abundance until 449.5 kyrs bp. Abundances then rose again until 447.5 kyrs bp, after which a second decline was observed, which culminated in the local extinction of the species between 440.0 and 439.1 kyrs bp. Based on the abundance, we could separate the *O. universa* population into two subsets (indicated as Intervals 1 and 2 respectively in Fig. 1) and treat them as a replication of the same general process.

*Trilobatus sacculifer* generally occurred at higher abundances of up to 38.4 % (on average 12.7 %) of the entire planktonic Foraminifera assemblage. From *c*.458 kyrs bp the abundance of the species decreased gradually until a local extinction between 439.1 and 438.6 kyrs bp, approximately 500–1000 yrs later than the comparable event in *O. universa*. Based on the species’ relative abundance we defined three phases of population size leading to the extinction (Phases 1–3 in Fig. 1). Phase 1 with high abundances (24 % mean) spans between 457.6 and 451.3 kyrs bp, Phase 2 with medium abundances (10 % mean) between 450.7 and 446.8 kyrs bp, and Phase 3 with low abundances (4 % mean) between 446.2 and 439.1 kyrs bp. Those phases could be explicitly investigated for morphological developments within the community.

#### 3.1.2 Morphotype abundances

In *O. universa*, two traditionally distinguished morphotypes (‘*B. bilobata*’ and ‘*O. suturalis*’) occurred with a mean abundance of 2.4 % (Fig. 1). ‘*Biorbulina bilobata*’ occurs in marginally higher abundances (on average 1.4 %) than ‘*O. suturalis*’ (on average 1.0 %). The abundance of abnormal morphotypes decreases over time, as shown by a GLM over the entire time period (*p <* 0.001, Nagelkerke-*R*^2^ = 0.22, Suppl. S1). When testing both intervals separately we observe that the decrease in abnormal morphotypes over time is significant for the second interval which leads to local extinction (*p* = 0.002, Nagelkerke-*R*^2^ = 0.22) but not for the first interval (*p* = 0.773, Nagelkerke-*R*^2^ = 0.01). In the *T. sacculifer* complex, we find higher abundances of the sacculifer-morphotype of up to 24 % broadly coinciding with Phase 1 of the abundance of the species (Fig. 1). During early Phase 2, this morphotype decreased rapidly in abundance and, although it never vanished completely within the limits of confidence, never comprised more than 10 % of the population from Phase 2 onwards. A constant decline over time is indicated by the GLM (*p <* 0.001, Nagelkerke-*R*^2^ = 0.56, Suppl. S1). This is confirmed by a *z*-test: The abundance of the sacculifer-morphotype differs significantly (*p <* 0.001) between all three Phases, with decreasing mean incidence of that morphotype from Phase 1 (14.7 %) over Phase 2 (5.8 %) to Phase 3 (2.0 %).

### 3.2 Morphology of *Orbulina universa* across replicated drops in abundance

Morphological parameters of *O. universa* are presented in Fig. 2. The shell size of *O. universa* specimens indicates the successive establishment of two populations, as indicated by Hartigan’s dip test and the coefficient of bimodality (Suppl. S1). Until 445.4 kyrs bp only one population with large shells (on average 399 µm) was present. After that, the size distribution reveals the existence of two populations with different sizes. One population shows an average size of 368 µm and can be considered a continuation of the population that was present before the split (Fig. 2a). While it differs in size from the larger population before the split on average (Mann–Whitney *U* test, *p <* 0.001) this can be explained by the gradual decrease in size of the larger population over time revealed by robust line fitting (*R*^2^ = 0.08, *p <* 0.001, Fig. 2a, Suppl. S1). The second population is significantly smaller (*p <* 0.001), only 155 µm on average, and only occurs with few specimens before the split from *c*.451 kyrs bp onwards. This population shows the same trend of decreasing size toward the local extinction (*R*^2^ = 0.12, *p <* 0.001, Fig. 2a, Suppl. S1). The large population became only marginally rarer; it was more abundant before the split, but never disappeared from the sedimentary record until the local extinction. Noteworthy, the coefficient of variation of the larger population was significantly dropping at the splitting point from 0.21 to 0.18. However, the smaller group showed an overall strongly reduced coefficient of variation of only 0.11 (Fig. 2b). The final split occurs within Interval 2 and is unrelated to the replicated abundance pattern in *O. universa*. Overall, shell size of the assemblage decreased toward the local extinction. When testing for a relationship between shell size and species abundance, a robust line fitting reveals no significant correlation between accumulation rates and shell size for the large (*p* = 0.070) population. For the small population, the regression is significant (*p <* 0.001) but explains very little variation in the data (*R*^2^ = 0.09, Suppl. S1).

**Figure 2.**
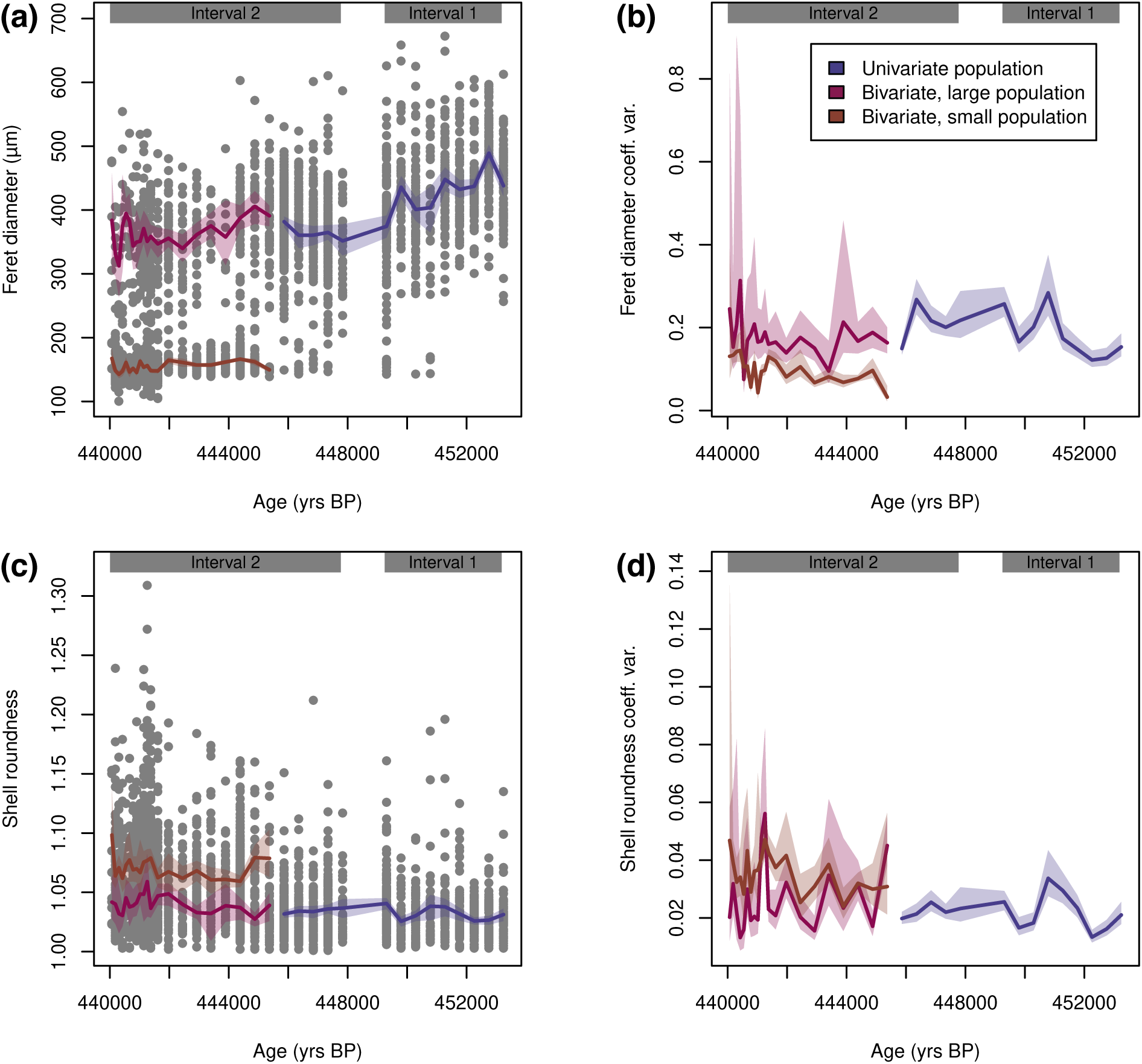
Morphology of *Orbulina universa* from marine isotope stage 12 in the Red Sea. (a) Shell size is showing bimodality after 445.4 kyrs BP, when a small population slowly appeared. (b) The variation in shell size is larger in the large population but generally decreased over time. (c) Shell roundness is rather stable in the large population, and considerably lower in the small population. (d) The variation in shell roundness is considerably higher in the small population but decreased slightly in the larger population toward local extinction. Raw values are plotted as grey dots, mean values as lines, and 95 % confidence intervals as shaded areas. The intervals based on species abundance (compare Fig. 1) are indicated.

To investigate shell roundness in *O. universa* we excluded ‘*B. bilobata*’ and ‘*O. suturalis*’ specimens (*n* = 59 specimens) from the analyses because both deviate significantly from the normal morphology of a terminal shell of that species. Shell roundness shows no indication of bimodality and is constantly decreasing over time according to a robust line fitting (*R*^2^ = 0.04, *p <* 0.001, Fig. 2c, Suppl. S1). The latter trend, however, is the result of the smaller population with inherently reduced shell roundness increasing in abundance. A secondary trend of shell roundness being correlated with species abundance exists, but it is much less pronounced (*R*^2^ =−0.01, *p <* 0.001, Suppl. S1). Shell roundness shows a decrease in variation in the large population toward the local extinction but is considerably higher in the small population (Fig. 2d). Noteworthy, the major trends in shell size and shell roundness are uncorrelated to the intervals defined by species abundance patterns.

#### 3.3 Morphology of *Trilobatus sacculifer* during a long, continuous extinction event

Morphological parameters of *T. sacculifer* are presented in Fig. 3. The shell size shows a comparable pattern to the incidence of the sacculifer-morphotype, with larger shells during Phase 1, a rapid size decrease during Phase 2, and small shells during Phase 3 (Fig. 3a). Comparing the values within the phases reveals a decrease in mean shell size from Phase 1 (411 µm) over Phase 2 (279 µm) to Phase 3 (237 µm). The differences in size are significant between all groups (*p <* 0.001 for a Kruskal–Wallis test, with all *p <* 0.001 in pairwise Mann–Whitney *U* tests). Moreover, the variation of shell size decreased significantly during Phase 2 (Fig. 3a). This decrease in shell size and variation is nearly exclusively caused by the lack of large specimens after Phase 2, while the size of the smallest specimens remained rather constant during the entire interval.

**Figure 3.**
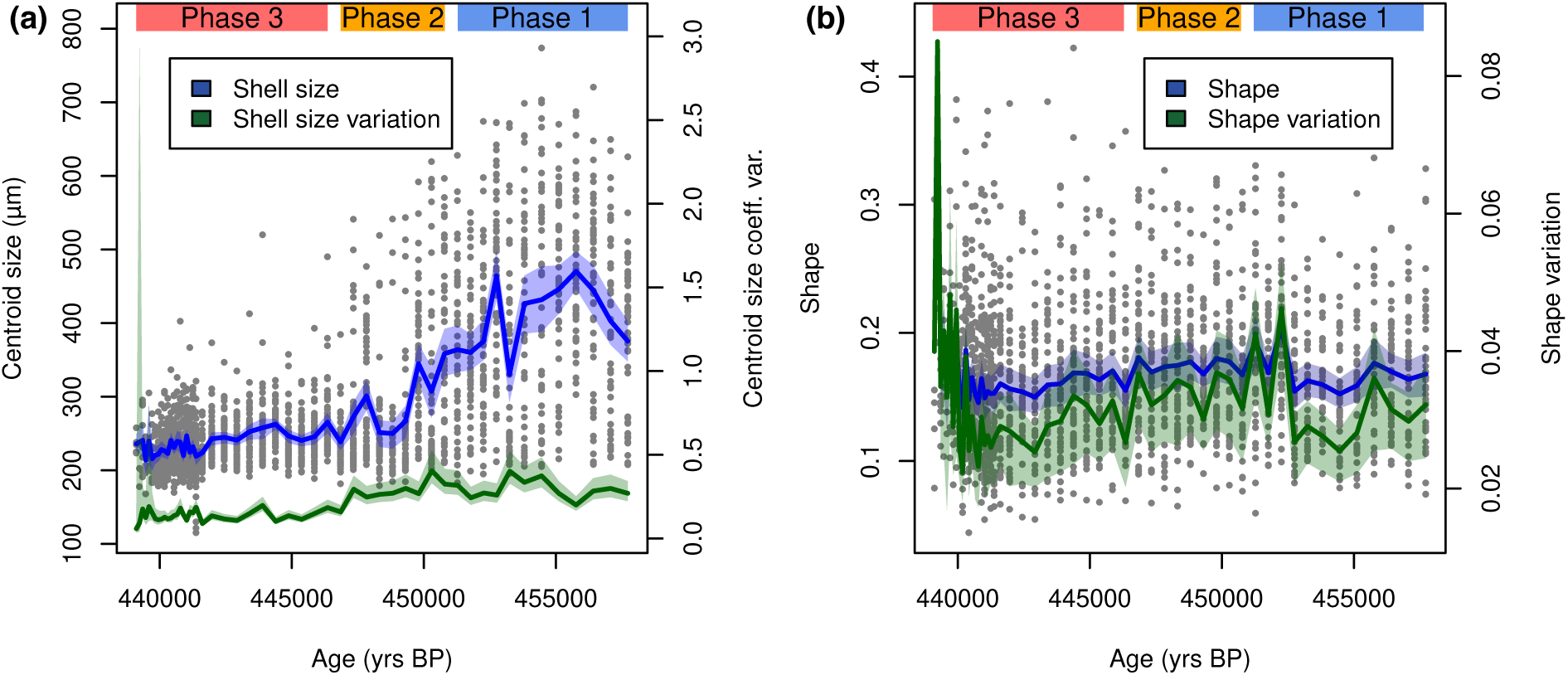
Morphology of *Trilobatus sacculifer* from marine isotope stage 12 in the Red Sea. The three phases defined by abundance (compare Fig. 1) are indicated. (a) Shell size and its variation decrease toward the local extinction (compare Suppl. S1). Raw values (grey dots) are plotted alongside the sample mean and coefficient of variation (solid lines) and their 95 % confidence intervals (shaded area). (b) Shell shape (Riemannian shape distance from the mean shape) excluding sacculifer-morphotypes (compare Suppl. S1). Raw values (grey dots) are plotted alongside the sample mean and standard deviation (solid lines) including the 95 % confidence interval (shaded area).

Using geometric morphometrics allows a more detailed analysis of the shape of *T. sacculifer* shells (Fig. 3b). Since the sacculifer-morphotype is a considerably derived morphology, we left it out of the dataset for most analyses (*n* = 136 specimens), but results including the sacculifer-morphotype specimens are presented in Suppl. S1 for comparison.

The shape of *T. sacculifer* specimens between phases differs significantly (NPMANOVA, *p <* 0.001). A *post-hoc* pairwise comparison of shape differences reveals, that the shape of *T. sacculifer* specimens differs between all three phases at *p* = 0.002, with a gradual shape change during the last *c*.10 000 yrs before local extinction (Fig. 3b). Shape variation showed no trend over time according to a robust line fitting (*R*^2^ = −0.1, *p* = 0.746), but would be significant when including the sacculifer-morphotype (*R*^2^ = 0.1, *p <* 0.001, Suppl. S1). This implies that a large part of the shape variation is linked to the disappearance of the sacculifer-morphotype in the population. Overall, however, there is a clear decreasing trend in shape variation between the end of Phase 1 and close to the end of Phase 3. The variation was rather low during early Phase 1 and seems to have increased very shortly before extinction, but the latter signal is accompanied by large confidence intervals due to the small sample size and should be treated with caution. This complex pattern likely obscures an overall trend, leading to an insignificant regression analysis. When comparing integrated values across the three phases defined by abundance patterns, the data reveal that variation was highest in Phase 2, mediocre in Phase 1, and lowest in Phase 3 (Fig. 4c). A *t*-test reveals that variation differences between Phases 1 and 2 are insignificant (*p* = 0.944), but specimens from Phase 3 differ significantly from specimens from both Phase 1 (*p* = 0.010) and Phase 2 (*p <* 0.001). This implies a significant reduction of phenotypic plasticity in the community when exposed to higher stress levels and impeding local extinction.

**Figure 4.**
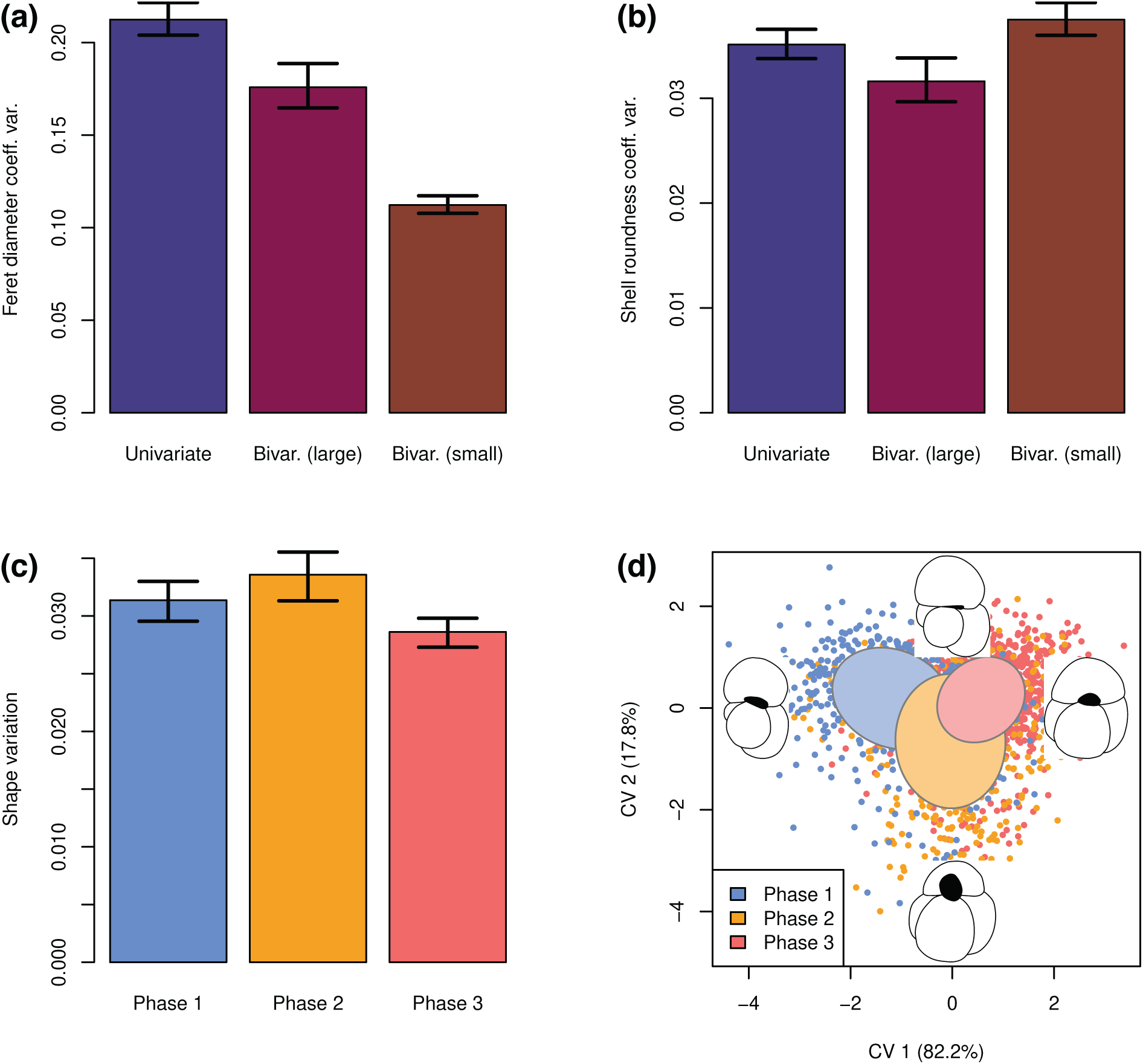
Stabilizing selection in planktonic Foraminifera during marine isotope stage 12 in the Red Sea. (a–b) The variation of shell size and shell roundness (including 95 % confidence interval) in the incumbent population of *Orbulina universa* decreased toward local extinction. The small population had an inherently reduced size variation but higher variation of shell roundness. (c) The shape variation (Riemannian shape distance from the mean shape, excluding sacculifer-morphotype) in *Trilobatus sacculifer* decreased significantly toward the local extinction (Phase 3, compare Fig. 1). Error bars depict the 95 % confidence interval. (d) Canonical variates analysis (CVA) of the shape of *T. sacculifer* (excluding sacculifer-morphotype). Points indicate specimens, ellipses indicate the 95 % confidence interval of the standard deviation on the centroid, black silhouettes depict the morphology at the extremal points of the canonical variate (CV) 1 and 2 axes, respectively.

When analysing the shape change from phase to phase (Fig. 4d) it is revealed that from Phase 1 to Phase 2 the terminal chamber became flatter and the aperture was inflated. At the same time the lower part of the shell containing the older chambers became more voluminous, so that there is a general trend towards a smaller terminal chamber in relation to the older chambers. This trend is reversed in Phase 3, with the terminal chamber becoming more inflated and the aperture flattening again. This led to a strong increase in the size of the terminal chamber in regard to the older shell.

## 4 Discussion

### 4.1 Error analysis

Morphometric data are especially prone to errors, because many things (like the orientation of specimens and digitization of landmarks) are done manually. We therefore analysed our data for the influence of potential error sources, following methods described by Yezerinac et al. (1992). We could not detect any errors of severe size in our data and show a full error discussion in Suppl. S1.

### 4.2 Morphological stabilization in planktonic Foraminifera

The concept of phenotypic plasticity (Schmalhauzen 1949) describes the observed morphological variation among specimens of a population, and is thus contrasted to variability, which is the potential of a population to vary within the borders of the genetic encoding (Wagner and Altenberg 1996). Importantly, phenotypic plasticity can only use pre-existing variability to adapt to environmental changes and is thus limited to a pre-adaptation of the population (Sultan and Stearns 2005). When faced with unprecedented environmental change, a population needs to innovate to overcome the new environmental stress by development of new mutations that serve adaptation (West-Eberhard 2003, Wagner 2011). This is especially important in planktonic Foraminifera, where the existence of (pseudo-)cryptic species can lead to an underestimation of genetic diversity and thus variability of the population. In *T. sacculifer* we can rule out that possibility, because despite its large morphospace it only contains one biological species with large variation (André et al. 2013). In *O. universa*, the morphospecies encompasses several biospecies (de Vargas et al. 1999), but at present only one genotype is known to occur in the Red Sea (Darling and Wade 2008).

Within our natural experiment, we observe bimodality in shell size coinciding with increasing salinity in the Red Sea in *O. universa*. A decrease in size and mean roundness of the incumbent large population is apparent when the species approached its local extinction in the aplanktonic zone. In both parameters, an unbiased random walk pattern best describes the observation (Table 1), implying that the observed morphological change is the result of accumulation of random changes rather than directional selection. The smaller population, however, shows stasis in both parameters without any distinctive trend one way or the other. A reduction in shell size has been proposed to indicate suboptimal environmental conditions (Ortiz et al. 1995, Schmidt et al. 2004) but also a buoyancy adaptation to increasing salinity in order to be able to stay at an optimum depth in denser water (Haenel 1987). Since the strong salinity increase in the Red Sea associated with the onset of the aplanktonic zone shifted the local habitat away from the optimum requirements of the incumbent species, it is therefore hard to say whether the trends we observe are the result of abiotic or biotic forcing. However, if the observed morphology change would be a purely biotic stress response, assuming that abundance is a useful proxy for the suitability of the environment (Drake and Griffen 2010), we would principally expect to see a comparable development in Interval 1 as in Interval 2. In contrast to that assumption, neither shell size nor shell roundness show any signs of a deviation associated with the first abundance drop at the end of Interval 1. We further observe that shell size and shell roundness are significantly different between both intervals at *p <* 0.001, which is not what one would expect if they would show a comparable internal pattern.

**Table 1.** | Results of phyletic evolution models of planktonic Foraminifera in the Red Sea during marine isotope stage 12.

| Parameter | Phyletic model | AIC<sub>c</sub> | Akaike weight |
| --- | --- | --- | --- |
| ***Orbulina universa*** |  |  |  |
| Shell size total | GRW | 377.84 | 0.306 |
|  | URW | 376.20 | 0.694 |
|  | Stasis | 425.81 | 0.000 |
| Shell size large pop. | GRW | 322.40 | 0.328 |
|  | URW | 320.96 | 0.672 |
|  | Stasis | 357.67 | 0.000 |
| Shell size small pop. | GRW | 157.96 | 0.014 |
|  | URW | 155.22 | 0.055 |
|  | Stasis | 149.58 | 0.931 |
| Shell roundness total | GRW | −246.40 | 0.371 |
|  | URW | −247.45 | 0.629 |
|  | Stasis | −192.50 | 0.000 |
| Shell round. large pop. | GRW | −257.97 | 0.200 |
|  | URW | −260.72 | 0.791 |
|  | Stasis | −251.90 | 0.010 |
| Shell round. small pop. | GRW | −131.90 | 0.034 |
|  | URW | −135.27 | 0.185 |
|  | Stasis | −138.16 | 0.781 |
| ***Trilobatus sacculifer*** |  |  |  |
| Shell size | GRW | 506.03 | 0.311 |
|  | URW | 504.44 | 0.689 |
|  | Stasis | 603.25 | 0.000 |
| Shell shape | GRW | −291.21 | 0.133 |
|  | URW | −293.50 | 0.416 |
|  | Stasis | −293.67 | 0.452 |

GRW: Generalized random walk, URW: Unbiased random walk, shell shape values of *T. sacculifer* excluding the sacculifer-morphotype

The observation of an incumbent large population which shows a morphological development characteristic for stress situations and the splitting of a small population showing stasis would be consistent with two possible endmember scenarios: (1) A monospecific *O. universa* population present at the time in the Red Sea shows signs of disruptive selection as a result of exposure to a suboptimal environment (Bull 1987), or (2) a new population comprising a different biospecies with different morphology is introduced into the Red Sea once environmental conditions become more suitable

To investigate those scenarios, we applied a Kendall rank-order correlation between individual shell size and shell roundness and found that they are significantly correlated (*ρ* = −0.321, *p <* 0.001, Suppl. S1). This implies that smaller individuals also tend to have less round shells. To test this, we divided the population into two subsets at the local minimum of the size distribution (238.6 µm) and tested the shell roundness in both subgroups against each other with a Mann–Whitney *U* test. The test confirms that the larger population produced shells that are on average significantly (*p <* 0.001) rounder (mean = 1.03) than the smaller subpopulation (mean = 1.07). The fact that the maximum shell roundness is practically 1.00 in both populations shows, that the observed trend is neither cause by a biological inability of shells to grow very round below a certain size nor the result of a lower precision of shell roundness determination in smaller shells. Comparing the 95 % confidence interval of the coefficient of variation shows a significantly higher variation of shell roundness in the population with smaller shells (0.036– 0.039) than in the population with larger shells (0.020–0.022, Fig. 4). Noteworthy, shell roundness variation did not meaningfully increase over time in either population, which shows the variation as inherent to the respective population (Fig. 2d). It is also possible that shell roundness and shell size are biologically integrated in *O. universa*, so that a change in one parameter necessitates a certain change in the other value (Klingenberg 2008). This would be an alternative explanation for the observed correlation between individual shell size and shell roundness, but the very low covariance between both parameters (−0.008) together with the fact that small specimens realize the complete range of possible deviations from sphericity principally argues against that hypothesis.

Our observations do not allow us to decide for one of the two potential hypotheses. The Red Sea was possibly invaded by a population with smaller, more variably shaped shells from approximately 445.4 kyrs bp onwards, which increasingly established its presence at the expense of the incumbent population due to their better adaptation to the local environment. Should this be true, we would find here the first example where different *O. universa* biospecies could be distinguished on the basis of relatively easily obtainable morphological characters (compare Morard et al. 2009) and also the first example of more than one *O. universa* biospecies occurring in the Red Sea (compare Darling and Wade 2008). Furthermore, it would provide evidence for different ecological preferences among those biospecies, which facilitates competitive exclusion due to increasing stress levels (Sultan and Stearns 2005). However, we would argue that this explanation is unlikely on the basis of other assumptions. (1) The potentially invading species would likely had had to be of Indian Ocean origin. Within an environmental setting of increasing salinity, it is hard to perceive that any species from the open-marine Indian Ocean would have a selective advantage over a native species from the Red Sea, that should be better adapted to high salinities (Sofianos and Johns 2003). (2) A trend of decreasing shell size with increasing salinity in *O. universa* was already observed by Haenel (1987), and was there attributed to buoyancy requirements, although Spero (1988) saw a closer relation to nutrient availability due to the energy requirements to build larger shells. This makes it reasonable that the observed size change is indeed an adaptive response by part of the population. (3) The splitting in two size populations as well as the increased variation in roundness in the smaller population both imply disrupt-ive selection (Badyaev 2005). We thus favour the explanation that the incumbent *O. universa* population underwent disruptive selection as a result of environmental stress, associated with drastically increasing salinity levels in the Red Sea. This would replicate results obtained on a Mediterranean population that was exposed to higher stress levels in relationship with the onset of sapropel deposition (Weinkauf et al. 2014), and would lend valuable evidence to the assumption, that certain morphological changes can be universally interpreted as indicator of environmental stress, at least when the same species are considered. The small subspecies would in that scenario be the better adapted result of accumulated random change (Hunt 2007), and would fall into a state of evolutionary stasis once apparently optimal adaptation is reached. This conforms with the proven development of stasis in extreme environments (Parsons 1994), while both populations can co-exist as long as an equilibrium point exists along the Lotka– Volterra isoclines (Sultan and Stearns 2005). Divergence without speciation is often observed in nature, and new character traits that are encompassed in the variability of the population can be fixed, once a diverging subpopulation occupies a new niche (West-Eberhard 2003). In the present case, the size difference in the two *O. universa* populations points toward a different in the depth habitat due to differential buoyancy (Haenel 1987). While thus disruptive selective pressure in *O. universa* may have led to the development of two populations, each individual population shows signs of stabilizing selection in its shell size parameter. It is very likely that the emerging new population is not the result of an evolutionary innovation, but the reactivation of pre-existing genetic variability in part of the population. This is evidenced by the rapid emergence via unbiased (in contrast to general) random walk and subsequent stasis of the small population as well as its ultimate failure to cope with the environmental change, which is not what would be expected under a scenario of dynamic evolution (Wagner 2011).

*Trilobatus sacculifer* draws a very similar picture in the form of decreasing variation in its shell size and shape over prolonged time spans before local extinction (Figs 3 and 4). In shell size this decrease complies with the unbiased random walk model, while for shell shape the distinction between unbiased random walk and stasis is nearly impossible (Table 1).

While the size decrease may itself be a signal for decreasing environmental suitability for the species (Ortiz et al. 1995, Schmidt et al. 2004), the decrease in variation is a clear signal for stabilizing selection inducing microenvironmental canalization (Waddington 1942, Schmalhauzen 1949). It is interesting, that stabilization occurs toward Phase 3, which must have been clearly environmentally suboptimal for *T. sacculifer* (drop in abundance, drastic increase in salinity). This could indicate either a stabilization of the regional environment, allowing selection for an optimal trait (Van Valen 1965), or a rapidly changing environment enforcing fluctuating selection that benefits a stable phenotype in the long run (Kawecki 2000, Pélabon et al. 2010). The slight increase of variation between Phases 1 and 2 may be an indicator for the second explanation, and indicate a Baldwin effect (Baldwin 1902), i.e. a phase of increased plasticity that allows adaptation to a new environment and evolutionary fixation afterwards. The reduction in the abundance of the sacculifer-morphotype after Phase 1 could be another manifestation of the stabilizing selection, but it could also have resulted from an inability of *T. sacculifer* shells to build an asymmetric chamber below a certain chamber size threshold. In the latter case, the drop in the abundance of the sacculifer-morphotype would result from the shell size decrease and not necessarily be a signal for stabilizing selection.

We observe certain morphological trends in *T. sacculifer* over time, as shown in Fig. 4, which can be interpreted as adaptive for a changing environment that gradually becomes less suitable for the population. During Phase 1, shells show more inflated and larger terminal chambers. This is reminiscent of the sacculifer-morphotype, which was left out of the CVA due to its strongly derived morphology. This phenotype is often assumed to be more abundant under lower stress levels (Caron et al. 1987a) but has already earlier been shown to be correlated with larger shell sizes (Hemleben et al. 1987). During Phase 2, coinciding with a first strong drop in sea level and thus salinity increase (Fig. 1), the abundance of the sacculifer-morphotype and the shell size decreased, while the size of the terminal chamber decreased as well, indicating a trend towards Kummerforms that are often associated with unfavourable environmental conditions (Berger 1969). Finally, in Phase 3 the shell size decreased further while the terminal chamber became larger again in relative terms. The latter trend could be necessitated by the small shell size, so that the terminal chamber had to become relatively larger again to provide enough space for gametogenesis. This phase is also characterized by a canalization peak, probably induced by an environment that was so unstable and unfavourable for the *T. sacculifer* population that any deviation from a very narrow morphotype at the edge of the species variability would drastically decrease survival rates and fitness. The entire investigated timespan can possibly be interpreted as follows: During Phase 1 an incumbent population of *T. sacculifer* was well adapted to the high but not excessive salinity levels in the Red Sea during MIS 12. During Phase 2, salinity levels started to rise, and induced a selection that via accumulation of random change led to a modification of the phenotype (Hunt 2007). In this phase, a Baldwin effect set in, increasing the variation of shape and thus buying the population time until further adaptation could settle in (Baldwin 1902). During Phase 3, the adaptation was as complete as it could possibly become within the variability of the species, and the successful morphologies became fixed via canalization (Schmalhauzen 1949, Gibson and Wagner 2000, West-Eberhard 2003). It is interesting to note that the trend towards smaller shells that was observed in *O. universa* is also present in *T. sacculifer*. This indicates that probably buoyancy problems due to the increasing water salinity were responsible for this adaptive trend.

*Trilobatus sacculifer* interestingly shows a noteworthy increase in shape variation very shortly before extinction, which to some degree seems to contradict the stabilizing trend that was prevalent until then. This trend could be interpreted in two ways. It was either a renewed, and in this case unsuccessful, Baldwin effect (Baldwin 1902). Alternatively, it may be the ultimate result of drastically increased fluctuating asymmetry within a population exposed to ever growing environmental change reaching the limits of its variability (Lens et al. 2002, Hendrickx et al. 2003). In both cases, it indicates a population under extensive stress levels and close to its collapse in absence of successful adaptation.

### 4.3 Planktonic foraminiferal adaptability

In our study we analysed the adaptive patterns of two species of planktonic Foraminifera that were exposed to increasing levels of salinity stress. Interestingly, the observations in both species coincide very well. Both species ultimately show signs of stabilizing selection in both shell morphology (Fig. 4) and the decreasing incidence of more derived phenotypes (Fig. 1). They already become extinct at water salinities which other protist groups still tolerate (Fenton et al. 2000, Zhang et al. 2010, Strom et al. 2013, Balzano et al. 2015), showing that the environmental change was not so strong that an adaptation toward it would be biologically infeasible. Both species showed processes that are well in line with adaptive patterns that use pre-existing variability to cope with environmental stress and did not seem able to innovate to overcome the stress levels (Waddington 1942, Schmalhauzen 1949, Waddington 1957, West-Eberhard 2003, Wagner 2011). This is despite the fact that planktonic Foraminifera have very short reproduction cycles of only 4–6 weeks (Bé 1977), which makes evolutionary processes fast enough that the time encompassed in this natural experiment would have been sufficient for new traits to emerge to adapt to the environmental change (Pawlowski et al. 1997).

It therefore seems that at least these two species of planktonic Foraminifera primarily adapt to changing environments through their innate variability, and show a low tendency toward innovation (i.e. a low evolvability, Wagner 2011). This means that these taxa would be heavily influenced by the exposition to environments outside their pre-adapted range and would prefer stress avoidance (i.e. migration) over innovative adaptation. If this pattern is prevalent amongst all planktonic Foraminifera, it could explain why the global assemblages faithfully tracked climate zone shifts during the Quaternary glacial–interglacial cycles (Prell and Damuth 1978, Thunell and Belyea 1982, Kučera 2007) rather than going through phases of intense radiation.

This study analysed only two species in a limited stress scenario, and further analyses are strictly required to confirm the reduced evolvability in planktonic Foraminifera, although other studies point toward similar conclusions (Weinkauf et al. 2014, Brombacher et al. 2017). If this lack of evolvability should be confirmed on a larger scale across planktonic Foraminifera however, it would make the entire group vulnerable to unprecedented environmental change scenarios. Given the capital role this taxon plays in marine carbonate sequestration and the carbon cycle (Schiebel 2002), this could be a major factor in future climate change scenarios. We therefore advocate for more work in this field to reach a better understanding of planktonic foraminiferal adaptive capabilities and modes of adaptation to environmental protrusions.

## Acknowledgements

We thank the members of the micropalaeontology workgroup at the MARUM Bremen and Minigraduiertenkolleg Tübingen for fruitful discussions concerning results of this study. Kerstin Braun (Arizona State University) is thanked for counting part of the foraminiferal assemblages used in this study. Katherine Grant (Australian National University) is recognized for providing the age–depth tie points for our age model. The principal author received funding from the Ministerium für Wissenschaft, Forschung und Kunst via the Landesgraduierten-Förderung (LGFG) and the Cushman Foundation for Foraminiferal Research via the Joanna M. Resig Fellowship.

## References

Akaike H (1974) A new look at the statistical model identification. IEEE Transactions on Automatic Control 19 (6): 716–23. doi:10.1109/TAC.1974.1100705.

Anderson MJ (2001) A new method for nonparametric multivariate analysis of variance. Austral Ecology 26 (1): 32–46. doi:10.1111/j.1442-9993.2001.01070.pp.x.

André A, Weiner A, Quillévéré F, Aurahs R, Morard R, Douady ChJ., de Garidel-Thoron Th, Escarguel G, de Vargas C, and Kucera M (2013) The cryptic and the apparent reversed: Lack of genetic differentiation within the morphologically diverse plexus of the planktonic foraminifer *Globigerinoides sacculifer*. Paleobiology 39 (1): 21–39. doi:10.1666/0094-8373-39.1.21.

Badyaev AV (2005) Role of stress in evolution: From individual adaptability to evolutionary adaptation. In: Hallgrímsson B and Hall BK (eds). Variation: A Central Concept in Biology. (Burlington, San Diego, and London: Elsevier). Chap. 13, pp. 277–302.

Baldwin JM (1902) Development and Evolution. (London: Macmillan & Co., Ltd.). 395 p.

Balzano S, Abs E, and Leterme SC (2015) Protist diversity along a salinity gradient in a coastal lagoon. Aquatic Microbial Ecology 74 (3): 263–77. doi:10.3354/ame01740.

Baumfalk YA, Troelstra SR, Ganssen G, and Van Zanen MJL (1987) Phenotypic variation of *Globorotalia scitula* (Foraminiferida) as a response to Pleistocene climatic fluctuations. Marine Geology 75: 231–40. doi:10.1016/0025-3227(87)90106-X.

Bé AWH (1977) An ecological, zoogeographic and taxonomic review of recent planktonic Foraminifera. In: Ramsay ATS (ed.). Oceanic Micropalaeontology. Vol. 1. (London, New York, and San Francisco: Academic Press). Chap. 1, pp. 1–100.

Benjamini Y and Yekutieli D (2001) The control of the false discovery rate in multiple testing under dependency. The Annals of Statistics 29 (4): 1165–88. doi:10.1214/aos/1013699998.

Berger WH (1969) Planktonic Foraminifera: Basic morphology and ecologic implications. Journal of Paleontology 43 (6): 1369–83.

Bookstein FL (1989) Principal warps: Thin-plate splines and the decomposition of deformations. IEEE Transactions on Pattern Analysis and Machine Intelligence 11 (6): 567–85. doi:10.1109/34.24792.

Brady HB (1877) Supplementary note on the Foraminifera of the Chalk(?) of the New Britain Group. Geological Magazine 4 (12): 535.

Brombacher A, Wilson PA, Bailey I, and Ezard ThHG (2017) The breakdown of static and evolutionary allometries during climatic upheaval. The American Naturalist 190 (3): 350–62. doi:10.1086/692570.

Bull JJ (1987) Evolution of phenotypic variance. Evolution: International Journal of Organic Evolution 41 (2): 303–15. http://www.jstor.org/stable/2409140.

Caron DA, Faber Jr. WW, and Bé AWH (1987a) Effects of temperature and salinity on the growth and survival of the planktonic foraminifer *Globigerinoides sacculifer*. Journal of the Marine Biological Association of the United Kingdom 67 (2):323–41. doi:10.1017/S0025315400026643.

Caron DA, Faber Jr. WW, and Bé AWH (1987b) Growth of the spinose planktonic foraminifer *Orbulina universa* in laboratory culture and the effect of temperature on life processes. Journal of the Marine Biological Association of the United Kingdom 67 (2): 343–58. doi:10.1017/S0025315400026655.

Claude J (2008) Morphometrics with R. Use R! (New York: Springer-Verlag). 316 p. doi:10.1007/978-0-387-77790-0.

Darling KF and Wade ChM (2008) The genetic diversity of planktic Foraminifera and the global distribution of ribosomal RNA genotypes. Marine Micropaleontology 67: 216–38. doi:10.1016/j.marmicro.2008.01.009.

Davison AC and Hinkley DV (1997) Bootstrap Methods and their Applications. Cambridge Series in Statistical and Probabilistic Mathematics 1. (Cambridge: Cambridge University Press). 582 p.

Drake JM and Griffen BD (2010) Early warning signals of extinction in deteriorating environments. Nature 467 (7314): 456–9. doi:10.1038/nature09389.

Ellison AM (1987) Effect of seed dimorphism on the density-dependent dynamics of experimental populations of *Atriplex triangularis* (chenopodiaceae). American Journal of Botany 74 (8): 1280– doi:10.1002/j.1537-2197.1987.tb08741.x.

Fenton M, Geiselhart S, Rohling EJ, and Hemleben Ch (2000) Aplanktonic zones in the Red Sea. Marine Micropaleontology 40 (3): 277–94. doi:10.1016/S0377-8398(00)00042-6.

Fisher RA (1936) The use of multiple measurements in taxonomic problems. Annals of Human Genetics 7 (2): 179–88. doi:10.1111/j.1469-1809.1936.tb02137.x.

Fligner MA and Killeen TJ (1976) Distribution-free two-sample tests for scale. Journal of the American Statistical Association 71 (353): 210–3. doi:10.1080/01621459.1976.10481517.

Furlow FB, Armijo-Prewitt T, Gangestad SW, and Thornhill R (1997) Fluctuating asymmetry and psychometric intelligence. Proceedings of the Royal Society B: Biological Sciences 264: 823–9. doi:10.1098/rspb.1997.0115.

Gibson G and Wagner G (2000) Canalization in evolutionary genetics: A stabilizing theory? BioEssays 22 (4): 372–80. doi:10.1002/(SICI)1521-1878(200004)22:4<372::AID-BIES7>3.0.CO;2-J.

Grant KM, Rohling EJ, Bronk Ramsey C, Cheng H, Edwards RL, Florindo F, Heslop D, Marra F, Roberts AP, Tamisiea ME, and Williams F (2014) Sea-level variability over five glacial cycles. Nature Communications 5: Article 5076. doi:10.1038/ncomms6076.

Haenel P (1987) Intérêt paléoocéanographique d’*Orbulina universa* d’Orbigny (foraminifère). Oceanologica Acta 10 (1): 15–25.

Hartigan JA and Hartigan PM (1985) The dip test of unimodality. The Annals of Statistics 13 (1): 70–84. doi:10.1214/aos/1176346577.

Hemleben Ch, Spindler M, Breitinger I, and Ott R (1987) Morphological and physiological responses of *Globigerinoides sacculifer* (Brady) under varying laboratory conditions. Marine Micropaleontology 12: 305–24. doi:10.1016/0377-8398(87)90025-9.

Hendrickx F, Maelfait J.-P, and Lens L (2003) Relationship between fluctuating asymmetry and fitness within and between stressed and unstressed populations of the wolf spider *Pirata piraticus*. Journal of Evolutionary Biology 16: 1270–9. doi:10.1046/j.1420-9101.2003.00633.x.

Hesemann M (2009) The Foraminifera.eu internet project. In: Peryt D and Kaminski MA (eds). 7th Micropalaeontological Workshop Abstracts and Excursion Guide: Mikro-2009, Sw. Katarzyna, Poland, 28–30 September 2009. Special Publications (Grzybowski Foundation), pp. 33–4. http://gf.tmsoc.org/Documents/Mikro2009/GFSP15.pdf.

Heslop D, De Schepper S, and Proske U (2011) Diagnosing the uncertainty of taxa relative abundances derived from count data. Marine Micropaleontology 79: 114–20. doi:10.1016/j.marmicro.2011.01.007.

Hunt G (2006) Fitting and comparing models of phyletic evolution: Random walks and beyond. Paleobiology 32 (4): 578–601. doi:10.1666/05070.1.

Hunt G (2007) The relative importance of directional change, random walks, and stasis in the evolution of fossil lineages. Proceedings of the National Academy of Sciences of the United States of America 104 (47): 18404–8. doi:10.1073/pnas.0704088104.

Kawecki TJ (2000) The evolution of genetic canalization under fluctuating selection. Evolution: International Journal of Organic Evolution 54 (1): 1–12. doi:10.1111/j.0014-3820.2000.tb00001.x.

Kendall DG (1984) Shape manifolds, Procrustean metrics, and complex projective spaces. Bulletin of the London Mathematical Society 16 (2): 81–121. doi:10.1112/blms/16.2.81.

Kendall MG (1938) A new measurement of rank correlation. Biometrika 30 (1–2): 81–93. doi:10.1093/biomet/30.1-2.81.

Klingenberg ChP (2003) A developmental perspective on developmental instability: Theory, models and mechanisms. In: Polak M (ed.). Developmental Instability: Causes and Consequences. (New York: Oxford University Press), pp. 14–34.

Klingenberg ChP (2008) Morphological integration and developmental modularity. Annual Review of Ecology, Evolution, and Systematics 39: 115–32. doi:10.1146/annurev.ecolsys.37.091305.110054.

Knappertsbusch M (2016) Evolutionary prospection in the Neogene planktic foraminifer *Globorotalia menardii* and related forms from ODP Hole 925B (Ceara Rise, western tropical Atlantic): Evidence for gradual evolution superimposed by long distance dispersal? Swiss Journal of Palaeontology 135 (2): 205–48. doi:10.1007/s13358-016-0113-6.

Kruskal WH and Wallis WA (1952) Use of ranks in one-criterion variance analysis. Journal of the American Statistical Association 47 (260): 583–621. doi:10.1080/01621459.1952.10483441.

Kucera M (2007) Planktonic Foraminifera as tracers of past oceanic environments. In: Hillaire-Marcel C, de Vernal A, and Chamley H (eds). Proxies in Late Cenozoic Paleoceanography. Developments in Marine Geology 1. (Amsterdam: Elsevier). Chap. 6, pp. 213–62. doi:10.1016/S1572-5480(07)01011-1.

Lens L, Van Dongen S, Kark S, and Matthysen E (2002) Fluctuating asymmetry as an indicator of fitness: Can we bridge the gap between studies? Biological Reviews of the Cambridge Philosophical Society 77: 27–38. doi:10.1017/S1464793101005796.

Malmgren BA (1984) Analysis of the environmental influence on the morphology of *Ammonia beccarii* (Linné) in southern European salinas. Geobios 17 (6): 737–46.

Malmgren BA and Kennett JP (1976) Biometric analysis of phenotypic variation in recent *Globigerina bulloides* d’Orbigny in the southern Indian Ocean. Marine Micropaleontology 1: 3–25. doi:10.1016/0377-8398(76)90003-7.

Mann HB and Whitney DR (1947) On a test of whether one of two random variables is stochastically larger than the other. The Annals of Mathematical Statistics 18 (1): 50–60. doi:10.1214/aoms/1177730491.

Marschner L, Staniek J, Schuster S, Triebskorn R, and Köhler H.-R (2013) External and internal shell formation in the ramshorn snail *Marisa cornuarietis* are extremes in a continuum of gradual variation in development. BMC Developmental Biology 13: Article 22. doi:10.1186/1471-213X-13-22.

Mary Y and Knappertsbusch MW (2013) Morphological variability of menardiform globorotalids in the Atlantic Ocean during Mid-Pliocene. Marine Micropaleontology 101: 180–93. doi:10.1016/j.marmicro.2012.12.001.

Meilland J, Siccha M, Weinkauf MFG, Jonkers L, Morard R, Baranowski U, Baumeister A, Bertlich J, Brummer G.-J, Debray P, Fritz-Endres Th, Groeneveld J, Magerl L, Munz P, Rillo MC, Schmidt Ch, Takagi H, Theara G, and Kucera M (2019) Highly replicated sampling reveals no diurnal vertical migration but stable species-specific vertical habitats in planktonic Foraminifera. Journal of Plankton Research 41 (2): 127–41. doi:10.1093/plankt/fbz002.

Moller T, Schulz H, and Kucera M (2013) The effect of sea surface properties on shell morphology and size of the planktonic foraminifer *Neogloboquadrina pachyderma* in the North Atlantic. Palaeogeography, Palaeoclimatology, Palaeoecology 391:34–48. doi:10.1016/j.palaeo.2011.08.014.

Morard R, Quillévéré F, Escarguel G, Ujiié Y, de Garidel-Thoron Th, Norris RD, and de Vargas C (2009) Morphological recognition of cryptic species in the planktonic foraminifer *Orbulina universa*. Marine Micropaleontology 71: 148–65. doi:10.1016/j.marmicro.2009.03.001.

Moritz C and Agudo R (2013) The future of species under climate change: Resilience or decline? Science 341 (6145): 504–8. doi:10.1126/science.1237190.

Nagelkerke NJD (1991) A note on a general definition of the coefficient of determination. Biometrika 78 (3): 691–2. doi:10.1093/biomet/78.3.691.

Nelder JA and Wedderburn RWM (1972) Generalized linear models. Journal of the Royal Statistical Society, Series A: General 135 (3): 370–84. http://www.jstor.org/stable/2344614.

Nellen W, Bettac W, Roether W, Schnack D, Thiel H, Weikert H, and Zeitschel B (1996) MINDIK Reise Nr. 5 [MINDIK Cruise No. 5]. Vol. 2. Meteor Berichte. (Hamburg: Leitstelle Meteor). 179 p. http://www.dfg-ozean.de/fileadmin/DFG/Berichte/M5b_Meteor_96-2.pdf.

D’Orbigny AD (1839) Foraminifères. In: de la Sagra R (ed.). Histoire physique et naturelle de l’Ile de Cuba. (Paris: A. Bertrand), p. 82.

Ortiz JD, Mix AC, and Collier RW (1995) Environmental control of living symbiotic and asymbiotic Foraminifera of the California Current. Paleoceanography 10 (6): 987–1009. doi:10.1029/95PA02088.

Osterauer R, Marschner L, Betz O, Gerberding M, Sawasdee B, Cloetens P, Haus N, Sures B, Triebskorn R, and Köhler H.-R (2010) Turning snails into slugs: Induced body plan changes and formation of an internal shell. Evolution & Development 12 (5): 474–83. doi:10.1111/j.1525-142X.2010.00433.x.

Parsons PA (1994) Morphological stasis: An energetic and ecological perspective incorporating stress. Journal of Theoretical Biology 171 (4): 409–14. doi:10.1006/jtbi.1994.1244.

Pawlowski J, Bolivar I, Fahrni JF, de Vargas C, Gouy M, and Zaninetti L (1997) Extreme differences in rates of molecular evolution of Foraminifera revealed by comparison of ribosomal DNA sequences and the fossil record. Molecular Biology and Evolution 14 (5): 498–505.

Peeters F, Ivanova E, Conan S, Brummer G.-J, Ganssen G, Troelstra S, and van Hinte J (1999) A size analysis of planktic Foraminifera from the Arabian Sea. Marine Micropaleontology 36 (1):31–63. doi:10.1016/S0377-8398(98)00026-7.

Pélabon Ch, Hansen ThF, Carter AJR, and Houle D (2010) Evolution of variation and variability under fluctuating, stabilizing, and disruptive selection. Evolution: International Journal of Organic Evolution 64 (7): 1912–25. doi:10.1111/j.1558-5646.2010.00979.x.

Prell WL and Damuth JE (1978) The climate-related diachronous disappearance of *Pulleniatina obliquiloculata* in late Quaternary sediments of the Atlantic and Caribbean. Marine Micropaleontology 3 (3): 267–77. doi:10.1016/0377-8398(78)90031-2.

Prell WL, Martin A, Cullen JL, and Trend M (1999a) The Brown University Foraminiferal Data Base. IGBP PAGES/World Data Center-A for Paleoclimatology Data Contribution Series 1999-027. Boulder: NOAA/NGDC Paleoclimatology Program. http://www.ncdc.noaa.gov/paleo/metadata/noaa-ocean-5908.html.

Prell WL, Martin A, Cullen JL, and Trend M (1999b) The Brown University Foraminiferal Data Base (BFD). http://doi.pangaea.de/10.1594/PANGAEA.96900.

R Core Team (2018) R: A Language and Environment for Statistical Computing. R Foundation for Statistical Computing. (Vienna). http://www.R-project.org/.

Rebotim A, Voelker AHL, Jonkers L, Waniek JJ, Meggers H, Schiebel R, Fraile I, Schulz M, and Kucera M (2017) Factors controlling the depth habitat of planktonic Foraminifera in the subtropical eastern North Atlantic. Biogeosciences 14: 827–59. doi:10.5194/bg-14-827-2017.

Rohling E, Abu-Zied R, Casford J, Hayes A, and Hoogakker B (2009) The marine environment: Present and past. In: Woodward J (ed.). The Physical Geography of the Mediterranean. The Oxford Regional Environments Series. (New York:Oxford University Press). Chap. 2, pp. 33–67.

Rostek F, Bard E, Beaufort L, Sonzogni C, and Ganssen G (1997) Sea surface temperature and productivity records for the past 240 kyr in the Arabian Sea. Deep-Sea Research, Part II: Topical Studies in Oceanography 44 (6–7): 1461–80. doi:10.1016/S0967-0645(97)00008-8.

Schiebel R (2002) Planktic foraminiferal sedimentation and the marine calcite budget. Global Biogeochemical Cycles 16 (4): Article 3. doi:10.1029/2001GB001459.

Schiebel R and Hemleben Ch (2017) Planktic Foraminifers in the Modern Ocean. (Berlin and Heidelberg: Springer-Verlag). 358 p. doi:10.1007/978-3-662-50297-6.

Schindelin J, Arganda-Carreras I, Frise E, Kaynig V, Longair M, Pietzsch T, Preibisch S, Rueden C, Saalfeld S, Schmid B, Tinevez J.-Y, White DJ, Hartenstein V, Eliceiri K, Tomancak P, and Cardona A (2012) Fiji: An open-source platform for biological-image analysis. Nature Methods 9: 676–82. doi:10.1038/nmeth.2019.

Schmalhauzen II (1949) Factors of Evolution: The Theory of Stabilizing Selection. (Madison: Blakiston Company). 327 p.

Schmidt DN, Renaud S, Bollmann J, Schiebel R, and Thierstein HR (2004) Size distribution of Holocene planktic foraminifer assemblages: Biogeography, ecology and adaptation. Marine Micropaleontology 50: 319–38. doi:10.1016/S0377-8398(03)00098-7.

Sen PK (1968) Estimates of the regression coefficient based on Kendall’s tau. Journal of the American Statistical Association 63 (324): 1379–89. http://www.jstor.org/stable/2285891.

Shapiro SS and Wilk MB (1965) An analysis of variance test for normality (complete samples). Biometrika 52 (3–4): 591–611. doi:10.1093/biomet/52.3-4.591.

Sofianos SS and Johns WE (2003) An Oceanic General Circulation Model (OGCM) investigation of the Red Sea circulation: 2. Three-dimensional circulation in the Red Sea. Journal of Geophysical Research, Oceans 108 (C3): Article 3066. doi:10.1029/2001JC001185.

Spero HJ (1988) Ultrastructural examination of chamber morphogenesis and biomineralization in the planktonic foraminifer *Orbulina universa*. Marine Biology 99 (1): 9–20. doi:10.1007/BF00644972.

Spezzaferri S, Kucera M, Pearson PN, Wade BS, Rappo S, Poole ChR, Morard R, and Stalder C (2015) Fossil and genetic evidence for the polyphyletic nature of the planktonic Foraminifera *“Globigerinoides”*, and description of the new genus *Trilobatus*. PLOS ONE 10 (5): e0128108. doi:10.1371/journal.pone.0128108.

Strom SL, Harvey EL, Fredrickson KA, and Mender-Deuer S (2013) Broad salinity tolerance as a refuge from predation in the harmful raphidophyte alga *Heterosigma akashiwo* (Raphidophyceae). Journal of Phycology 49 (1): 20–31. doi:10.1111/jpy.12013.

Sultan SE and Stearns SC (2005) Environmentally contingent variation: Phenotypic plasticity and norms of reaction. In: Hallgrímsson B and Hall BK (eds). Variation: A Central Concept in Biology. (Burlington, San Diego, and London: Elsevier). Chap. 14, pp. 303–32.

Theil H (1950) A rank-invariant method of linear and polynomial regression analysis, iii. Proceedings of the Koninklijke Nederlandse Akademie van Wetenschappen 53 (9): 1397–412.

Thunell RC, Locke SM, and Williams DF (1988) Glacio-eustatic sea-level control on Red Sea salinity. Nature 334: 601–4. doi:10.1038/334601a0.

Thunell R and Belyea P (1982) Neogene planktonic foraminiferal biogeography of the Atlantic Ocean. Micropaleontology 28 (4): 381–98. doi:10.2307/1485451.

Van Valen L (1965) Morphological variation and width of ecological niche. The American Naturalist 99 (908): 377–90. http://www.jstor.org/stable/2459179.

Vangel MG (1996) Confidence intervals for a normal coefficient of variation. The American Statistician 50 (1): 21–6. http://www.jstor.org/stable/2685039.

de Vargas C, Norris R, Zaninetti L, Gibb SW, and Pawlowski J (1999) Molecular evidence of cryptic speciation in planktonic foraminifers and their relation to oceanic provinces. Proceedings of the National Academy of Sciences of the United States of America 96: 2864–8. doi:10.1073/pnas.96.6.2864.

Venables WN and Ripley BD (2002) Modern Applied Statistics with S. 4th ed. Statistics and Computing. (Springer-Verlag). 495 p.

Waddington CH (1942) Canalization of development and the inheritance of aquired characters. Nature 150 (3811): 563–5. doi:10.1038/150563a0.

Waddington CH (1957) The Strategy of the Genes: A Discussion of some Aspects of Theoretical Biology. (London: Allen & Unwin). 262 p.

Wagenmakers E.-J and Farrell S (2004) AIC model selection using Akaike weights. Psychonomic Bulletin & Review 11 (1): 192–6. doi:10.3758/BF03206482.

Wagner A (2011) The Origins of Evolutionary Innovation: A Theory of Transformative Change in Living Systems. (New York: Oxford University Press). 253 p.

Wagner GP and Altenberg L (1996) Perspective: Complex adaptations and the evolution of evolvability. Evolution: International Journal of Organic Evolution 50 (3): 967–76. doi:10.2307/2410639.

Weinkauf MFG, Moller T, Koch MC, and Kucera M (2014) Disruptive selection and bet-hedging in planktonic Foraminifera: Shell morphology as predictor of extinctions. Frontiers in Ecology and Evolution 2: Article 64. doi:10.3389/fevo.2014.00064.

West-Eberhard MJ (2003) Developmental Plasticity and Evolution. (New York: Oxford University Press). 794 p.

Yezerinac SM, Lougheed SC, and Handford P (1992) Measurement error and morphometric studies: Statistical power and observer experience. Systematic Biology 41 (4): 471–82. doi:10.1093/sysbio/41.4.471.

Zelditch ML, Swiderski DL, and Sheets HD (2012) Geometric Morphometrics for Biologists: A Primer. 2nd ed. (London, Waltham, and San Diego: Academic Press). 478 p. http://booksite.elsevier.com/9780123869036/index.php.

Zhang J, Li Y, and Chen J (2010) Salinity tolerance and genetic diversity of the dinoflagellate *Oxyrrhis marina*. Journal of Ocean University of China 9 (1): 87–93. doi:10.1007/s11802-010-0087-8.

