## Supplementary material for "Extinctions in the marine plankton preceded by stabilizing selection": Suppl. S1

### *Electronic supplements to:* 'Extinctions in the marine plankton preceded by stabilizing selection'
bioRxiv

*Manuel F. G. Weinkauf*[1,2,3], *Fabian Bonitz*[1,a], *Rossana Martini*[3], *and Michal Kučera*[2]

[1]Department of Geosciences, Eberhard–Karls Universität Tübingen, Tübingen, Germany; [2]MARUM, Universität Bremen, Bremen, Germany; [3]Department of Earth Sciences, Université de Genève, Genève, Switzerland; [a]*Current address:* Norwegian Research Centre, Bjerknes Centre for Climate Research, Bergen, Norway

### 1 Material and methods details

#### 1.1 Morphological data extraction

**Table S1.** Description of landmarks used for *Trilobatus sacculifer* specimens from marine isotope stage 12 in the Red Sea in apertural standard orientation (compare Fig. S1) and their associated landmark type after Bookstein (1991).

| Landmark | Description | Type |
|---|---|---|
| 1 | Leftmost point of aperture | III |
| 2 | Topmost point of aperture in middle part (point of maximum curvature) | II |
| 3 | Rightmost point of aperture | III |
| 4 | Trisection between aperture, second-youngest, and third-youngest chamber | I |
| 5 | Lowermost point of aperture in middle part | III |
| 6 | Left intersection between youngest chamber and older shell | I |
| 7 | Left point of maximum curvature of youngest chamber | II |
| 8 | Topmost point of youngest chamber | III |
| 9 | Right point of maximum curvature of youngest chamber | II |
| 10 | Right intersection between youngest chamber and older shell | I |
| 11 | Intersection between third-oldest chamber and older shell | I |
| 12 | Intersection between second-oldest and third-oldest chamber | I |

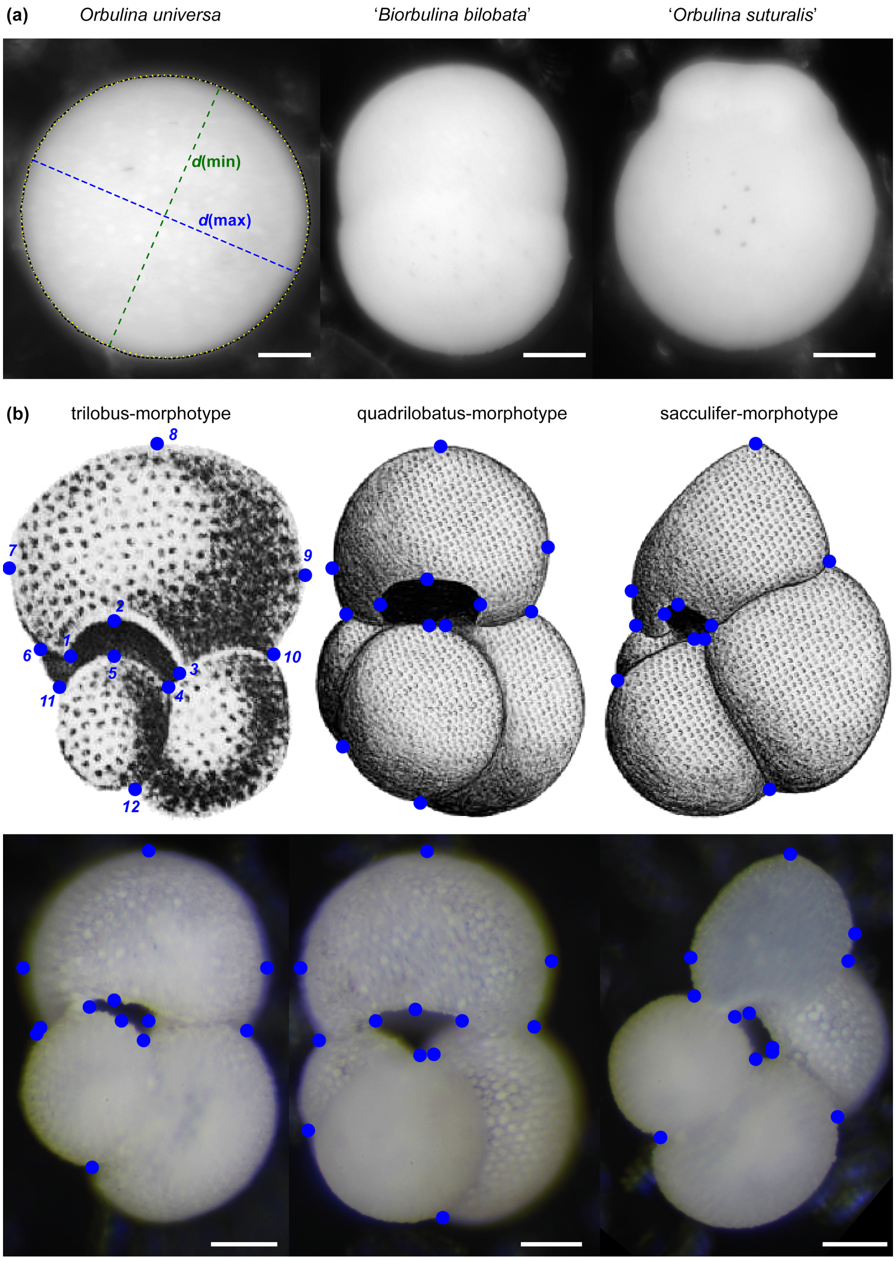


**Figure S1.** Depiction of species and measurements. (a) Measurements and morphotypes in *Orbulina universa*. Shell size was extracted as Feret diameter (not shown) according to the raw outline of the shell (black, dashed line). Shell roundness was calculated as ratio between the longest ($d$(max)) and shortest ($d$(min)) axis of an ellipse fitted to the outline (yellow, dotted line). The incidence of the ecophenotypes '*Biorbulina bilobata*' and '*Orbulina suturalis*' was counted manually. (b) In *Trilobatus sacculifer*, three morphotypes were distinguished, which are shown here in apertural standard orientation. Twelve landmarks were extracted for morphometric analyses (indicated as blue dots and exemplarily numbered in the drawing of the trilobus-morphotype, compare Table S1). The upper row shows type specimen drawings of the three morphotypes: trilobus-morphotype from Reuss (1850), quadrilobatus- and sacculifer-morphotypes from Banner and Blow (1960). The lower row shows corresponding light microscopy photographs from the studied samples. Scale bars for all light microscopy images equal 100 µm.

#### 2 Additional results

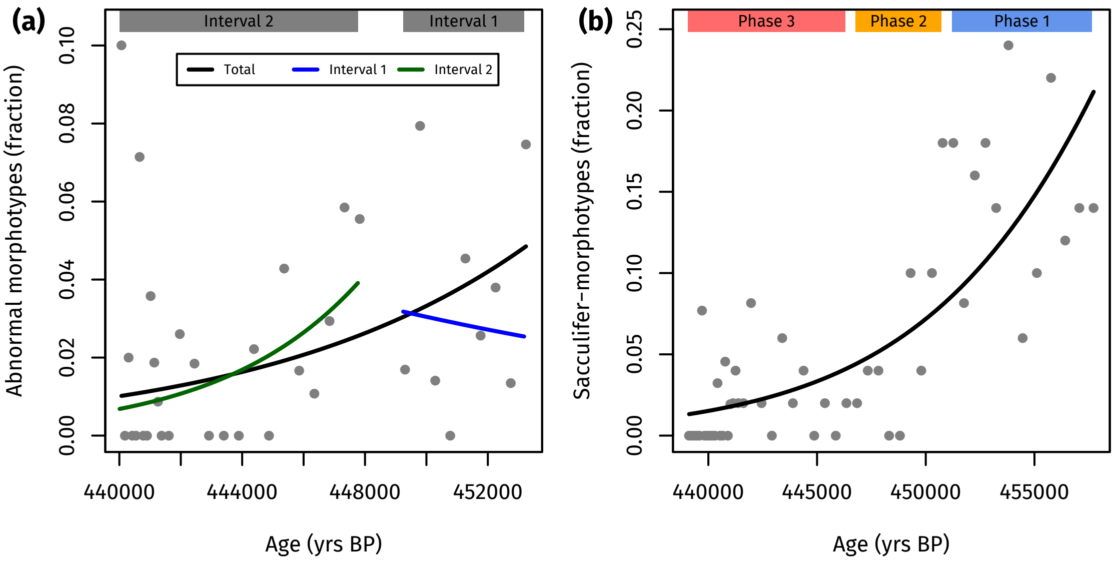


| | *O. universa* (total) | *O. universa* (Interval 1) | *O. universa* (Interval 2) | *T. sacculifer* |
|---|---|---|---|---|
| Nagelkerke-$R^2$ | 0.22 | 0.01 | 0.22 | 0.56 |
| $p$-value | <0.001 | 0.773 | 0.002 | <0.001 |

**Figure S2.** Generalized linear models (Nelder and Wedderburn 1972, binomial distribution, logit as link-function), of (a) the incidence of abnormal morphotypes in *Orbulina universa* and (b) the incidence of sacculifer-morphotypes in *Trilobatus sacculifer* from marine isotope stage 12 in the Red Sea. The intervals for *O. universa* and phases for *T. sacculifer* based on abundance patterns are indicated.

**Table S2.** Test for bimodality in shell size and shell roundness of *Orbulina universa* from marine isotope stage 12 in the Red Sea. Shown is the coefficient of bimodality after Ellison (1987) and *p*-values of Hartigan's dip test (Hartigan and Hartigan 1985) (compare Fig. S3).

| | | Shell size | | Shell roundness | |
|---|---|---|---|---|---|
| Age (yrs BP) | Depth in core (cm) | Coeff. bimodality | *p*-value | Coeff. bimodality | *p*-value |
| 440 068.01 | 1436.25 | 1.78 | 0.503 | 1.41 | 0.442 |
| 440 186.99 | 1436.75 | 0.85 | 0.915 | 0.54 | 0.749 |
| 440 305.97 | 1437.25 | 0.8 | 0.981 | 0.48 | 0.946 |
| 440 424.95 | 1437.75 | 0.85 | 0.528 | 0.49 | 0.546 |
| 440 543.94 | 1438.25 | 2.57 | 0.034 | 0.78 | 0.966 |
| 440 662.92 | 1438.75 | 0.88 | 0.216 | 0.51 | 0.204 |
| 440 781.9 | 1439.25 | 0.89 | 0.795 | 0.54 | 0.59 |
| 440 900.88 | 1439.75 | 0.85 | 0.819 | 0.46 | 0.48 |
| 441 019.86 | 1440.25 | 0.52 | 0 | 0.64 | 0.634 |
| 441 138.84 | 1440.75 | 0.86 | 0.492 | 0.51 | 0.566 |
| 441 257.82 | 1441.25 | 0.82 | 0.966 | 0.55 | 0.987 |
| 441 376.81 | 1441.75 | 0.87 | 0.383 | 0.56 | 0.689 |
| 441 614.77 | 1442.75 | 0.84 | 0.969 | 0.57 | 0.678 |
| 441 971.71 | 1444.25 | 0.68 | 0 | 0.51 | 0.722 |
| 442 447.64 | 1446.25 | 0.72 | 0.09 | 0.48 | 0.041 |
| 442 923.56 | 1448.25 | 0.88 | 0.018 | 0.45 | 0.789 |
| 443 401.5 | 1450.25 | 0.94 | 0.355 | 0.63 | 0.974 |
| 443 893.51 | 1452.25 | 0.85 | 0.955 | 0.37 | 0.921 |
| 444 385.52 | 1454.25 | 0.86 | 0.087 | 0.53 | 0.387 |
| 444 877.53 | 1456.25 | 0.82 | 0.009 | 0.52 | 0.607 |
| 445 369.54 | 1458.25 | 0.62 | 0.04 | 0.63 | 0.209 |
| 445 861.55 | 1460.25 | 0.35 | 0.994 | 0.41 | 0.846 |
| 446 353.56 | 1462.25 | 0.65 | 0.179 | 0.55 | 0.676 |
| 446 845.57 | 1464.25 | 0.31 | 0.818 | 0.52 | 0.471 |
| 447 337.58 | 1466.25 | 0.25 | 0.917 | 0.5 | 0.559 |
| 447 829.59 | 1468.25 | 0.33 | 0.931 | 0.59 | 0.964 |
| 449 305.62 | 1474.25 | 0.4 | 0.603 | 0.54 | 0.273 |
| 449 797.63 | 1476.25 | 0.34 | 0.944 | 0.55 | 0.989 |
| 450 289.64 | 1478.25 | 0.24 | 0.885 | 0.47 | 0.903 |
| 450 781.65 | 1480.25 | 0.55 | 0.938 | 0.74 | 0.847 |
| 451 273.66 | 1482.25 | 0.33 | 0.964 | 0.68 | 0.544 |
| 451 765.67 | 1484.25 | 0.45 | 0.944 | 0.62 | 0.835 |
| 452 257.68 | 1486.25 | 0.31 | 0.414 | 0.47 | 0.846 |
| 452 749.69 | 1488.25 | 0.41 | 0.944 | 0.58 | 0.505 |
| 453 241.69 | 1490.25 | 0.33 | 0.771 | 0.55 | 0.981 |

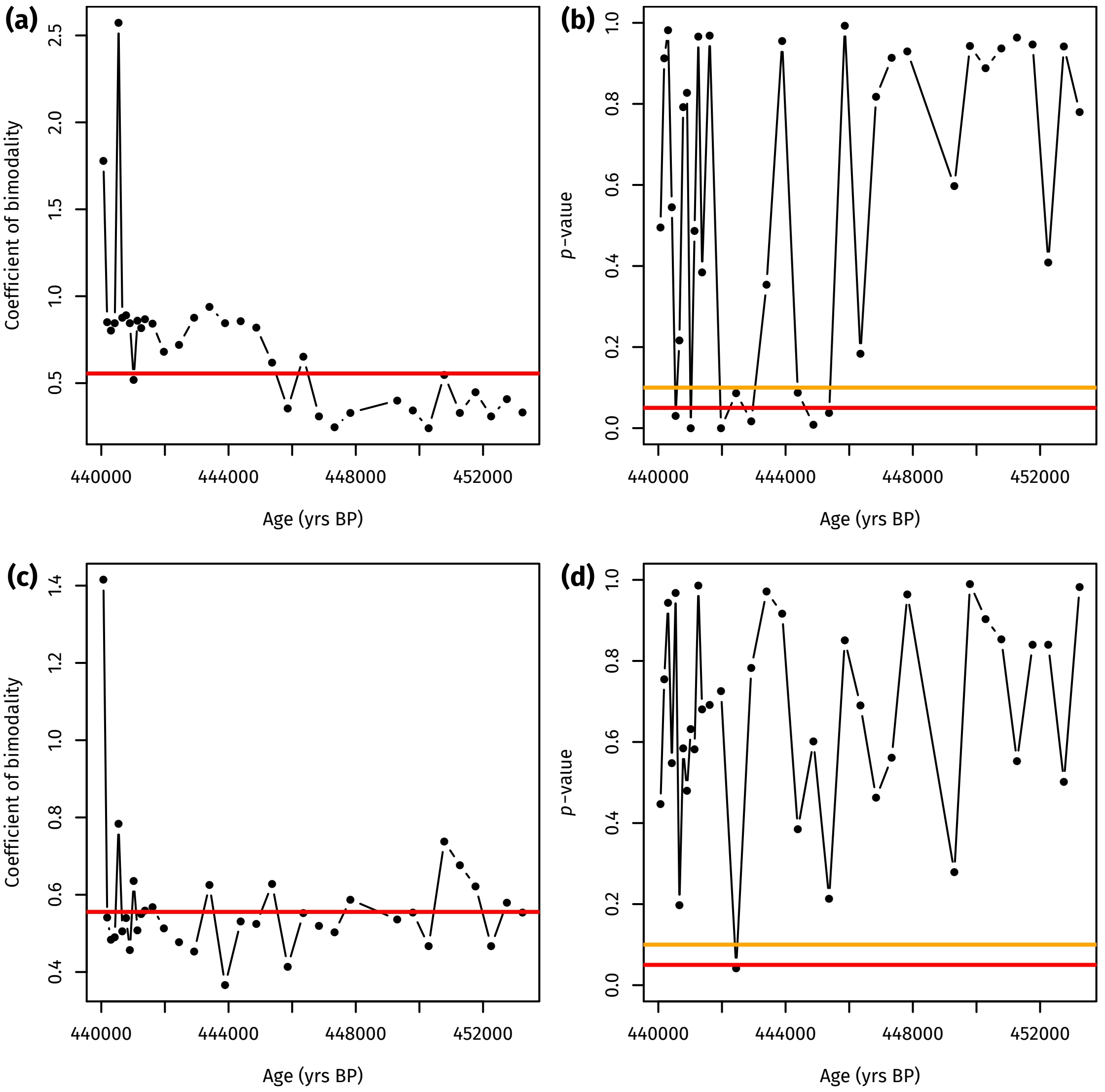


**Figure S3.** Test for bimodality in shell size (a, b) and shell roundness (c, d) of *Orbulina universa* from marine isotope stage 12 in the Red Sea. Shown is the coefficient of bimodality after Ellison (1987) (a, c), where a value larger $\frac{5}{9}$ (red line) indicates bimodality, and results of Hartigan's dip test (Hartigan and Hartigan 1985) (b, d), with indications of the $q = 0.05$ (red line) and $q = 0.10$ (orange line) confidence level. Both metrics indicate persistent bimodality in the shell size of the *O. universa* assemblage after 445.4 kyrs BP, but none in shell roundness values (compare Table S2).

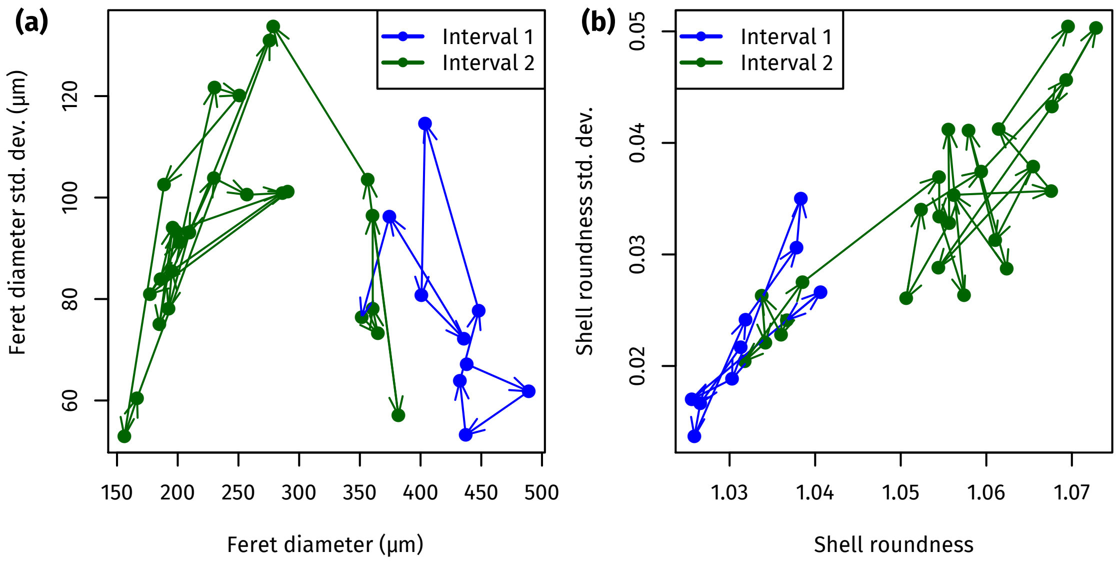


**Figure S4.** Morphology of *Orbulina universa* from marine isotope stage 12 in the Red Sea. Cross-plots of the entire population, where mean values per sample (points) are connected by arrows in temporal order. (a) Shell size decreases over time but shell size variation remains rather constant. (b) Shell roundness variation increases due to the increase in abundance of the more variable small population.

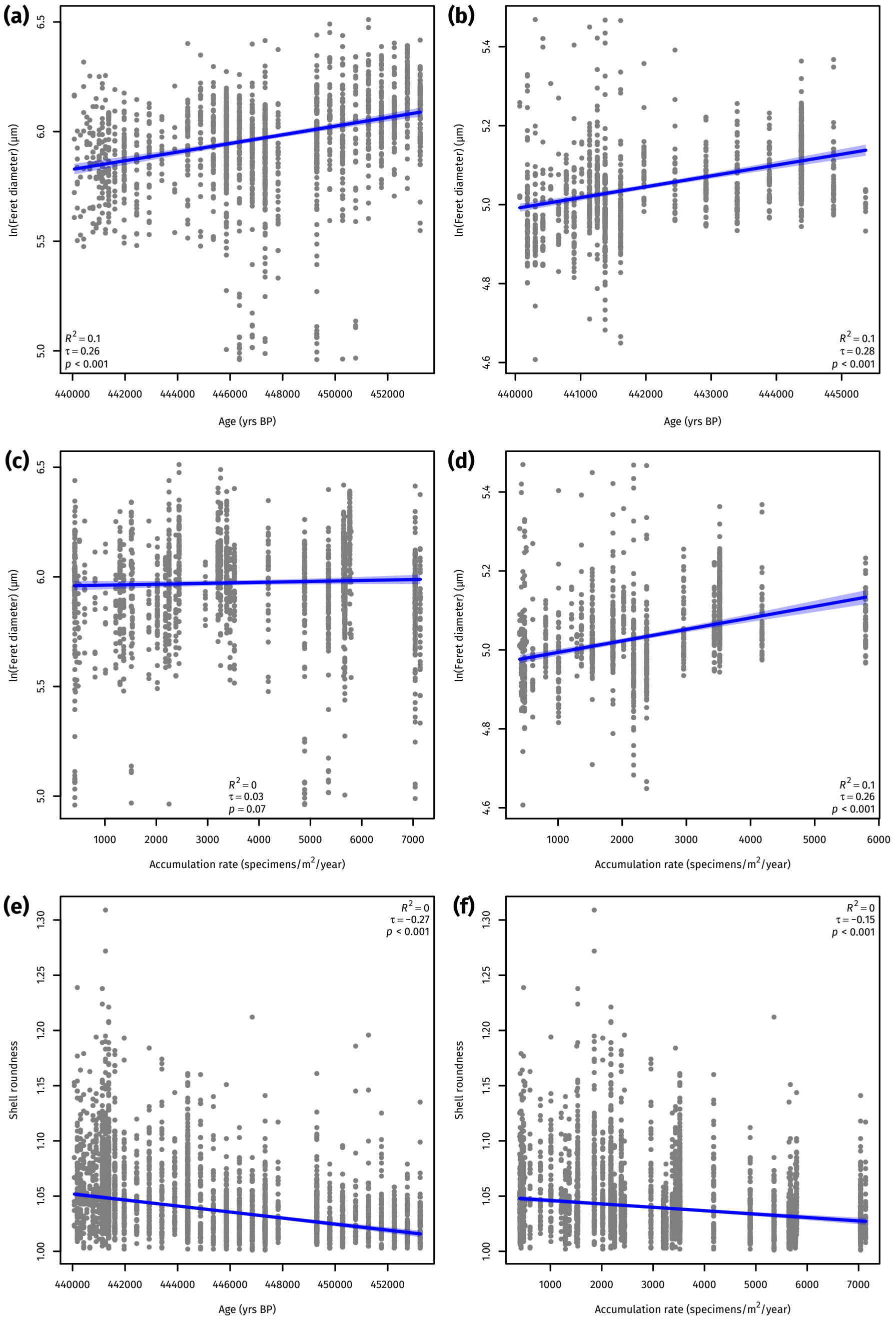


**Figure S5.** Shell size and roundness trends over time and with species abundance in *Orbulina universa* from marine isotope stage 12 in the Red Sea, using Kendall–Theil robust line fitting (Kendall 1938, Theil 1950, Sen 1968). (a, c) Shell size for the large population, that is present all the time. (b, d) Shell size for the small population, that appears at 445.4 kyrs BP. (e, f) Shell roundness for the entire population. The 95 % confidence interval of the regression is shown as shaded area. Shell sizes were $\log_e$-transformed prior to analysis.

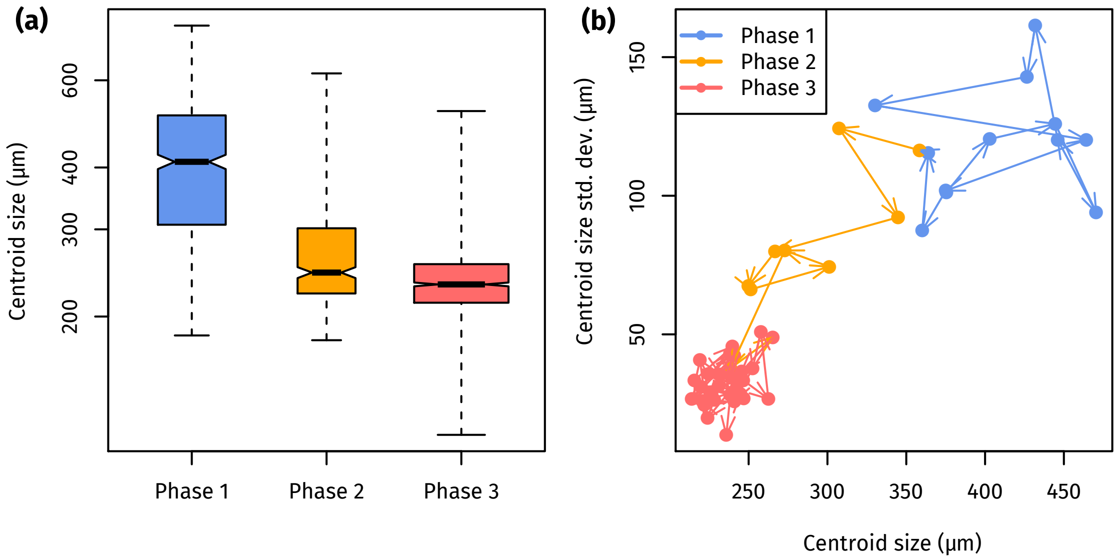


**Figure S6.** Shell morphology of *Trilobatus sacculifer* through three phases defined by species abundance from marine isotope stage 12 in the Red Sea. (a) Boxplot of shell sizes. Box width is scaled to the number of observations per group, the thick line indicates the median, the boxes extend to the interquartile range, the whiskers cover the entire range of observations. Note the log-scaling of the $y$-axis. (b) Cross-plot, where mean values per sample (points) are connected by arrows in temporal order.

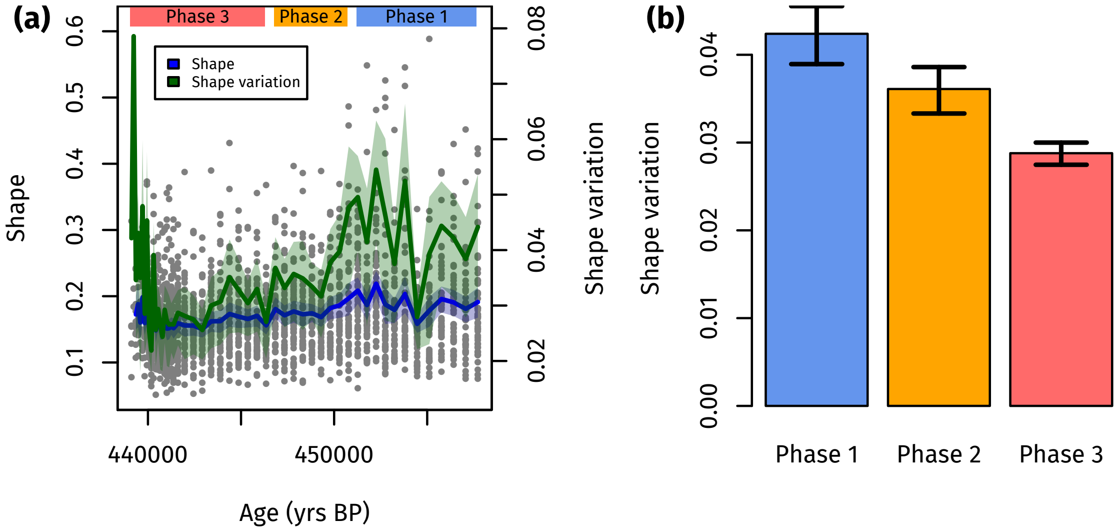


**Figure S7.** Shape of *Trilobatus sacculifer* from marine isotope stage 12 in the Red Sea, including sacculifer-morphotypes. (a) The shape (Riemannian shape distance) of the shells changed toward the local extinction and its variation dropped. Raw values (grey dots) are plotted alongside the sample mean and variation (solid lines). The three phases defined by abundance are indicated. (b) The shape variation (variance of the Riemannian shape distance from the grand mean) significantly decreased from Phase 1 to Phase 3. The shape between abundance groups differs significantly (NPMANOVA, Anderson 2001, $p < 0.001$), with all pairwise differences being significant ($p = 0.002$).

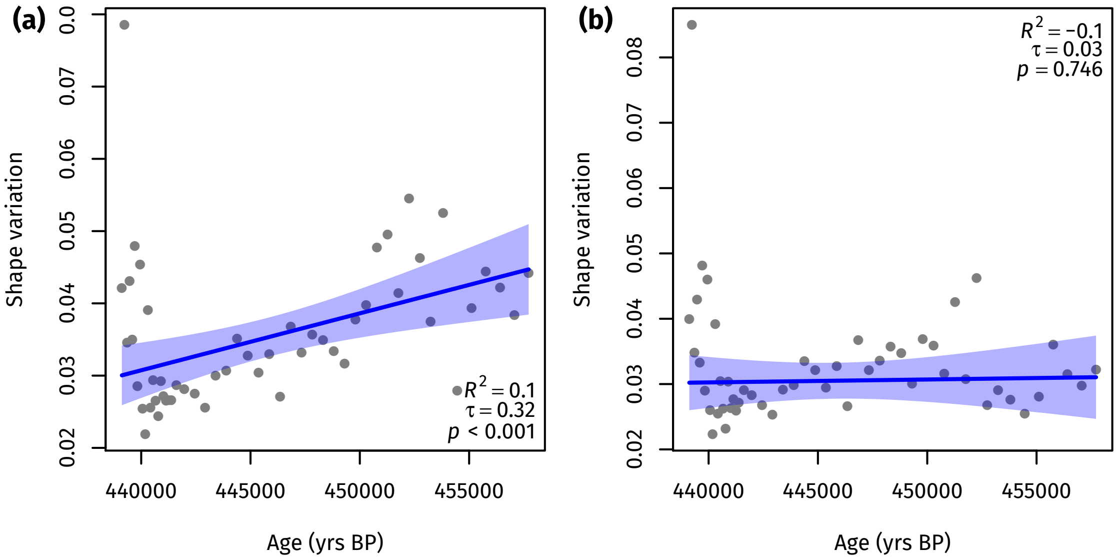


**Figure S8.** Shape variation of *Trilobatus sacculifer* from marine isotope stage 12 in the Red Sea, using Kendall–Theil robust line fitting (Kendall 1938, Theil 1950, Sen 1968). (a) When including the sacculifer-morphotype, the shape variation decreased over time. (b) This seems to be mainly related to the extinction of the sacculifer-morphotype, because this trend disappears when excluding this morphotype. The 95 % confidence interval of the regression is shown as shaded area.

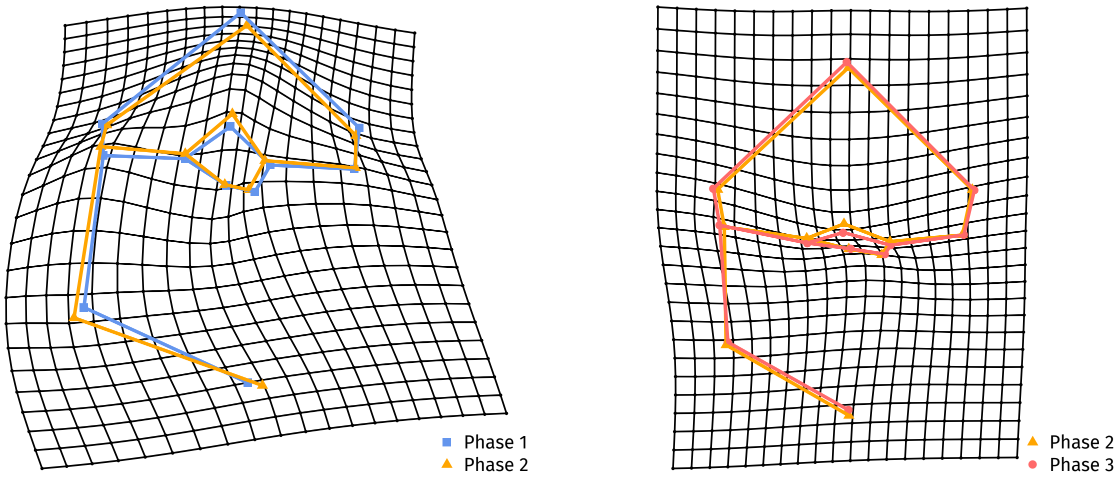


**Figure S9.** Thin plate splines of the deformation of *Trilobatus sacculifer* specimens from marine isotope stage 12 in the Red Sea between three phases defined by relative abundances.

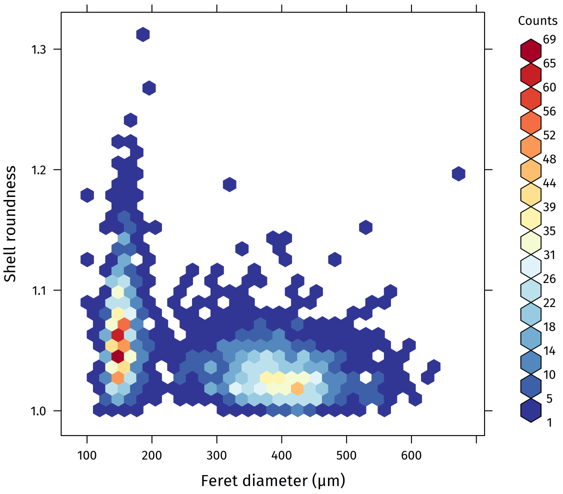


**Figure S10.** Correlation between shell size (as Feret diameter) and shell roundness in *Orbulina universa* during marine isotope stage 12 in the Red Sea. The population shows clear bimodality in shell size. Smaller specimens show a significantly higher variation of roundness than larger specimens.

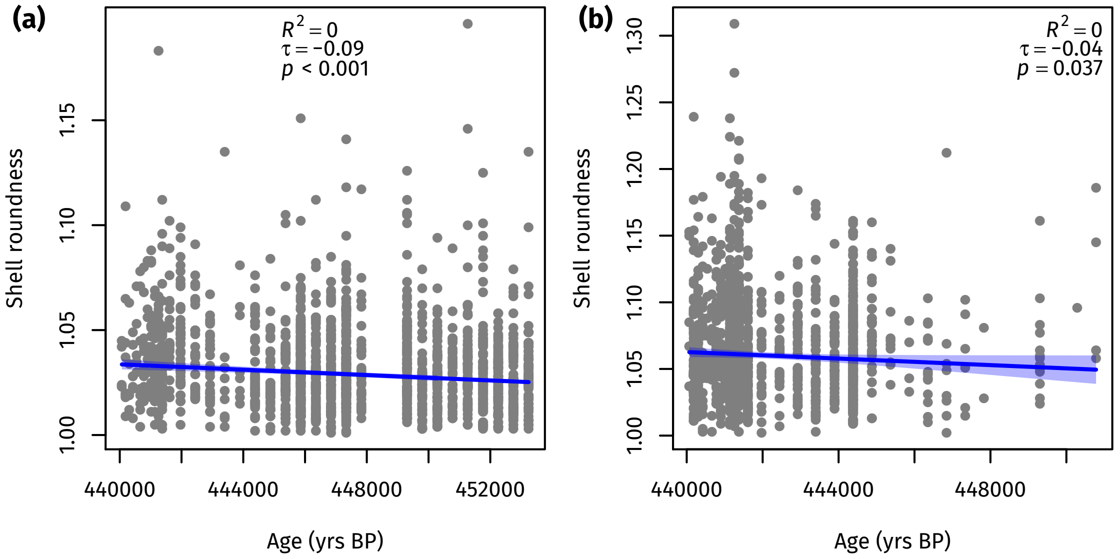


**Figure S11.** Shell roundness trend in the large (a) and small (b) population of *Orbulina universa* from marine isotope stage 12 in the Red Sea. The regression is significant for both populations but explains practically no variation in the data. The 95 % confidence interval of the regression is shown as shaded area.

### 3 Error discussion

Geometric morphometric analyses are error prone due to the large amount of manual steps involved, which can bias the reproducibility of the results. Amongst others, this includes the error of the measurement device and the measurer. We limited the device error by using the same microscope and camera for all photographs and using a constant magnification per species. The error by the measurer was limited by ensuring that all tasks were applied by the same researcher per specimen. Two other sources of error are only relevant for the geometric morphometric analyses of *Trilobatus sacculifer*. They were quantified using analysis of variances-based (ANOVA) approaches proposed by Yezerinac et al. (1992), and are discussed in the following.

#### 3.1 Error due to manual orientation of specimens

While all specimens per species were homogeneously oriented by the same researcher, eliminating a personal error term, a mis-orientation of specimens could still occur. To estimate that error, we repeatedly (eight times) reoriented three randomly selected specimens covering the whole observed size range, and estimated the landmark position error due to orientation. The analyses shows that manually orienting the specimens did not introduce a large error. The mean squares of the replicates (2219) is much smaller than the residual mean squares (23 739), with the ANOVA being highly insignificant ($p = 0.999$). This indicates a high reproducibility of landmarks regardless of small orientation errors, and the relative measurement error associated with specimen orientation sums up to only 1.07 %.

#### 3.2 Error due to manual landmark placement

A second source of error is the problem of misplacing landmarks in the specimen images. Due to the large amount of samples it was not feasible to replicate landmark extraction for all samples, instead we replicated this step for two samples, one with on average very small specimens (1439.5–1440 cm) and one with on average very large specimens (1488–1488.5 cm). The reasoning behind this is that larger specimens have a higher effective resolution, because they contain more pixels under constant magnification, and morphological details are better visible in larger specimens when the magnification is kept constant. It is thus likely that the landmark extraction error is not independent of specimen size. For both samples we replicated the landmark extraction and calculated the session error (i.e. the average mismatch between replicates) and the individual error (i.e. the error per specimen) as well as the relative measurement error (Table S3). In both samples we observe that the residual mean squares (error variance) of the session is much larger than the session factor mean squares, and that the ANOVA results for the session error are insignificant, indicating a high replicability of landmark positions in different sessions. The individual error ANOVA is highly significant in both cases, with the individual factor mean squares (between-specimen variance) being much larger than the residual mean squares (within-specimen variance). This indicates that the observed differences between specimens are much larger than what could be explained by errors in landmark extraction. Accordingly, the relative measurement errors associated with landmark misplacement are very small, with 0.264 % for small specimens and 0.148 % for large specimens.

**Table S3.** Error calculation for landmark extraction in *Trilobtaus sacculifer* specimens from marine isotope stage 12 in the Red Sea, exemplarily performed on two samples with on average very small (1439.5–1440 cm) and very large (1488–1488.5 cm) specimens.

| | | 1439.5–1440 cm | 1488–1488.5 cm |
|---|---|---|---|
| Session error | Session factor mean squares | 52 | 20 |
| | Residual mean squares | 4105 | 21194 |
| | *p*-value | 0.910 | 0.976 |
| Individual error | Individual factor mean squares | 8199 | 42357 |
| | Residual mean squares | 11 | 31 |
| | *p*-value | $< 0.001$ | $< 0.001$ |

#### 3.3 Allometry analyses

One last potential problem in morphometric analyses, especially in geometric morphometrics, can be the influence of ontogeny on shape. This can be tested in two ways (Zelditch et al. 2012). (1) One can adapt the univariate allometric equation and calculate the Riemannian shape distances between the smallest individual and all other individuals, and then calculate the regression between size and shape distance to the smallest individual. When doing this for our data for *T. sacculifer* (Fig. S12a) we find that the regression is insignificant ($p = 0.919$) and shell size explains less of the observed shape change than the null-model ($R^2 < -0.001$). (2) Alternatively, one can regress the partial warps of the landmark configuration on centroid size data in a multivariate regression approach to investigate the shape change as a whole dependent on size (Zelditch et al. 2012, eq. 11.4). Doing so reveals a significant allometric component in *T. sacculifer* shape ($p < 0.001$), but it explains only 4.10 % of the observed shape changes and is thus practically negligible, because it cannot be responsible for the signals we see in the data. It is furthermore mainly limited to a widening of the aperture (Fig. S12b), which is no trend that we observe in the *T. sacculifer* population in association with increasing stress levels. We thus conclude that allometry is no problem in our analysis of *T. sacculifer* shape changes with environmental stress.

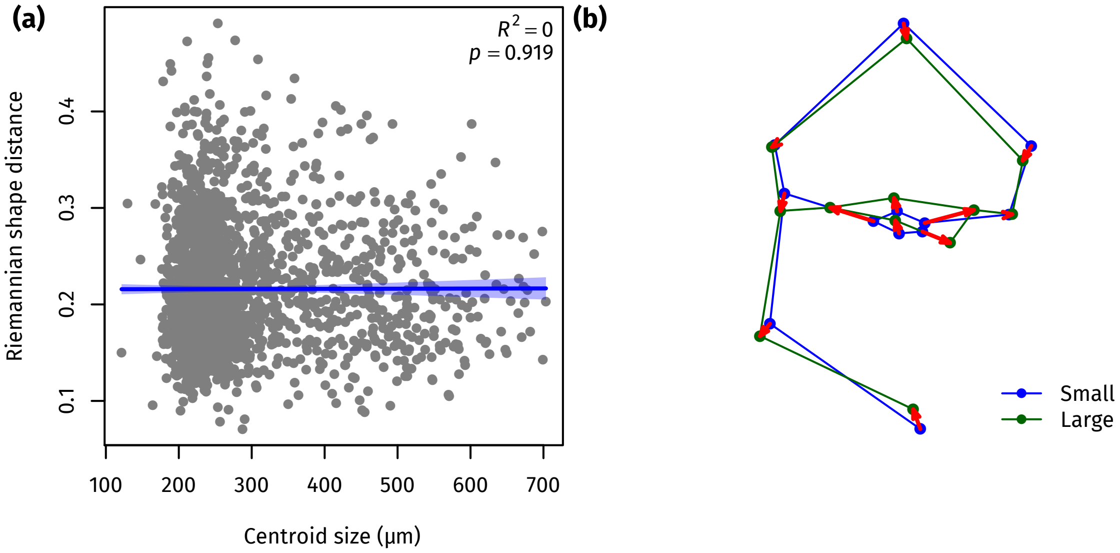


**Figure S12.** Allometry of *Trilobatus sacculifer* from marine isotope stage 12 in the Red Sea. (a) The univariate allometry approach shows no significant correlation between shape (expressed as Riemannian shape distance from the smallest individual) and size. (b) The shape change with ontogeny is nearly exclusively limited to a widening of the aperture.

### References


Anderson MJ (2001) A new method for non-parametric multivariate analysis of variance. *Austral Ecology* 26 (1): 32–46. DOI: 10.1111/j.1442-9993.2001.01070.pp.x.

Banner FT and Blow WH (1960) Some primary types of species belonging to the superfamily Globigerinaceae. *Contributions from the Cushman Foundation for Foraminiferal Research* 11 (1): 1–41.

Bookstein FL (1991) *Morphometric Tools for Landmark Data: Geometry and Biology*. (Cambridge, New York, and Melbourne: Cambridge University Press). 435 p.

Ellison AM (1987) Effect of seed dimorphism on the density-dependent dynamics of experimental populations of *Atriplex triangularis* (chenopodiaceae). *American Journal of Botany* 74 (8): 1280–8. DOI: 10.1002/j.1537-2197.1987.tb08741.x.

Hartigan JA and Hartigan PM (1985) The dip test of unimodality. *The Annals of Statistics* 13 (1): 70–84. DOI: 10.1214/aos/1176346577.

Kendall MG (1938) A new measurement of rank correlation. *Biometrika* 30 (1–2): 81–93. DOI: 10.1093/biomet/30.1-2.81.

Nelder JA and Wedderburn RWM (1972) Generalized linear models. *Journal of the Royal Statistical Society, Series A: General* 135 (3): 370–84. URL: http://www.jstor.org/stable/2344614.

Reuss AE (1850) Neue Foraminiferen aus den Schichten des österreichischen Tertiärbeckens. *Denkschriften der Kaiserlichen Akademie der Wissenschaften, Mathematisch–Naturwissenschaftliche Classe* 1: 365–90.

Sen PK (1968) Estimates of the regression coefficient based on Kendall's tau. *Journal of the American Statistical Association* 63 (324): 1379–89. URL: http://www.jstor.org/stable/2285891.

Theil H (1950) A rank-invariant method of linear and polynomial regression analysis, iii. *Proceedings of the Koninklijke Nederlandse Akademie van Wetenschappen* 53 (9): 1397–412.

Yezerinac SM, Lougheed SC, and Handford P (1992) Measurement error and morphometric studies: Statistical power and observer experience. *Systematic Biology* 41 (4): 471–82. DOI: 10.1093/sysbio/41.4.471.

Zelditch ML, Swiderski DL, and Sheets HD (2012) *Geometric Morphometrics for Biologists: A Primer*. 2nd ed. (London, Waltham, and San Diego: Academic Press). 478 p. URL: http://booksite.elsevier.com/9780123869036/index.php.
